# A multi-scale pipeline linking drug transcriptomics with pharmacokinetics predicts *in vivo* interactions of tuberculosis drugs

**DOI:** 10.1101/2020.09.03.281550

**Authors:** Joseph Cicchese, Awanti Sambarey, Denise Kirschner, Jennifer Linderman, Sriram Chandrasekaran

## Abstract

Tuberculosis (TB) is the deadliest infectious disease worldwide. The design of new treatments for TB is hindered by the large number of candidate drugs, drug combinations, dosing choices, and complex pharmaco-kinetics/dynamics (PK/PD). Here we study the interplay of these factors in designing combination therapies by linking a machine-learning model, INDIGO-MTB, which predicts *in vitro* drug interactions using drug transcriptomics, with a multi-scale model of drug PK/PD and pathogen-immune interactions called *GranSim*. We calculate an *in vivo* drug interaction score (iDIS) from dynamics of drug diffusion, spatial distribution, and activity within lesions against various pathogen sub-populations. The iDIS of drug regimens evaluated against non-replicating bacteria significantly correlates with efficacy metrics from clinical trials. Our approach identifies mechanisms that can amplify synergistic or mitigate antagonistic drug interactions *in vivo* by modulating the relative distribution of drugs. Our mechanistic framework enables efficient evaluation of *in vivo* drug interactions and optimization of combination therapies.

## Introduction

Tuberculosis (TB), caused by inhalation of *Mycobacterium tuberculosis* (Mtb), remains the world’s deadliest infectious disease, infecting 30% of all people world-wide and leading to ~1.3 million deaths annually^1,2^. The emergence of multidrug resistance coupled with slow progress in developing new drugs has created a pressing need to identify new approaches to treat TB. The current standard TB treatment regimen is a combination of 4 first-line anti-TB drugs – the antibiotics isoniazid (H), rifampicin (R), pyrazinamide (Z), and ethambutol (E). This treatment has remained unchanged over 50 years^3–5^. Typical combination therapy for TB is administered for at least six months, while treating drug-resistant strains may take up to two years. New drugs, such as bedaquiline, linezolid, and pretomanid, are being tested in new regimens to potentially shorten TB treatment^6,7^ The WHO has called for entirely new strategies to meet the goals for ‘End TB’, which aims to reduce TB deaths by 95% by 2035.

The large number of potential drug combinations greatly complicates TB treatment design^8^. Therapy involving drug combinations can lead to surprising non-linear effects; some drugs can enhance each other’s action leading to higher potency (synergy), or drugs can interfere with their action leading to reduced potency (antagonism)^9,10^. These drug interactions can impact treatment efficacy and emergence of drug resistance^11^. Such synergistic or antagonistic drug interactions can be determined using checkerboard assays by screening a panel of drug combinations in multiple doses against Mtb^12^. However, such experimental screening of drug interactions has limited throughput despite recent developments in reducing the number of doses required for measurement^13–15^. Designing an optimal 4-drug combination from a set of just 50 candidate drugs at a single dose requires ~200,000 drug interaction experiments. The dosage and dosing frequency further increase the space of possible regimens exponentially^16^.

Measuring *in vivo* drug interactions is even more challenging as it requires mice, primates, or other model organisms infected with Mtb^17^. Consequently, the number of drug candidates that can be screened through these model organisms is very limited. Further, current drug screening strategies for TB do not consider a patient’s immune system. Once Mtb is inhaled, it triggers a cascade of immune responses that result in the accumulation of an immune cell-rich mass around infected cells and bacteria known as a granuloma. Mtb can persist for decades within granulomas, and there are multiple granulomas within lungs of infected patients^18^. Granulomas also create a physical barrier altering the penetration of drugs, which can greatly impact relative drug concentrations at the site of infection and lead to effective mono-therapies in granulomas^19–22^. Granulomas also produce nutrient-starved and hypoxic environments that contain Mtb that are phenotypically tolerant to antibiotics, further complicating treatment^23,24^.

This study addresses these challenges by creating a multi-scale pipeline combining two cutting-edge computational approaches, operating at different biological scales, to evaluate combination therapies using drug transcriptomics and pharmacokinetics /pharmacodynamics (PK/PD) (Figure 1). First, to rapidly predict drug-drug interactions (synergy/antagonism) among combinations of two or more drugs, we utilize the existing *in silico* tool—<u>in</u>ferring <u>d</u>rug <u>i</u>nteractions using chemo<u>g</u>enomics and <u>o</u>rthology (INDIGO) optimized for Mtb (INDIGO-MTB)^8,12^. INDIGO-MTB uses a training data of known drug interactions along with drug transcriptomics data as inputs. INDIGO-MTB then utilizes a machine-learning algorithm to identify gene expression patterns that are predictive of specific drug-drug interactions. Once trained, INDIGO-MTB can determine if new drugs in combination have synergistic or antagonistic interactions using transcriptomics data. We previously used INDIGO-MTB to identify synergistic drug regimens for treating TB from over a million possible drug combinations using the pathogen response transcriptome elicited by individual drugs. The INDIGO-MTB model contains 164 drugs with anti-TB activity and it accurately predicted novel interactions of two-drug and three-drug combinations *in vitro*^8^.

**Figure 1:**
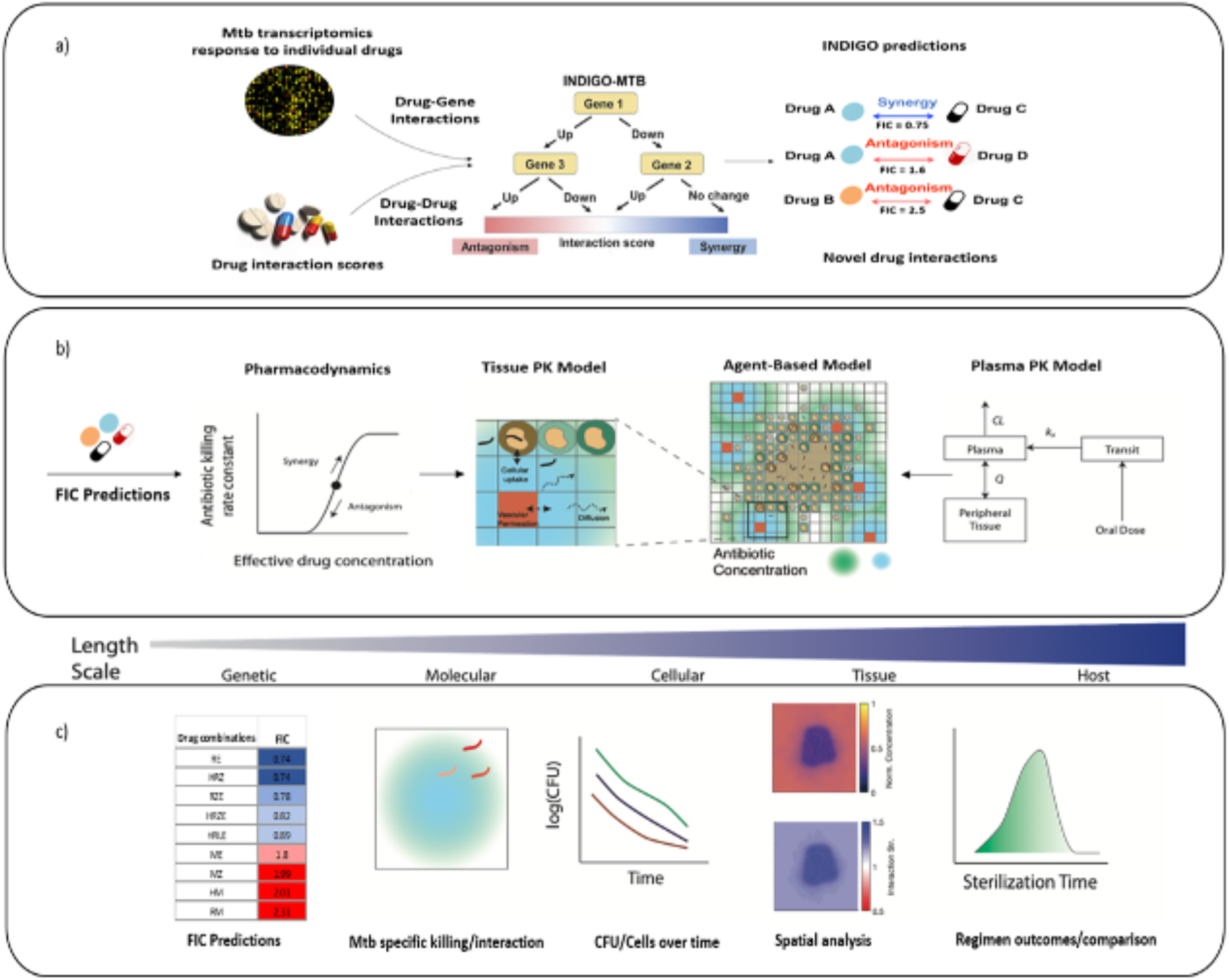
Overview of our multiscale pipeline to predict *in vivo* drug interactions. a) INDIGO-MTB uses Mtb transcriptomic responses to drugs and experimentally measured drug-drug interactions as inputs for training a machine-learning model, inferring synergistic and antagonistic interactions between new drug combinations as output. b) Components of the model integrating GranSim and INDIGO-MTB. From right to left, the plasma PK model determines the time-dependent concentration of all antibiotics following oral doses, which in turn determines the amount of antibiotic delivered onto the agent-based model grid. The computational grid is 200×200 square grid spaces, representing 4mm by 4mm of lung. Within the agent-based model, the tissue PK model describes antibiotic diffusion and binding as well as immune cell accumulation. Based on the local concentration of antibiotics, the PD model evaluates an antibiotic killing rate constant based on an effective concentration that is calculated from each individual antibiotic concentration. The corresponding FIC predicted from INDIGO-MTB either increases or decreases this effective concentration, depending on whether the combination is synergistic or antagonistic. c) Different predictions and outcomes, with the gradient above corresponding to the relevant length scale for the model/prediction. From left to right, predictions made by integration of GranSim and INDIGO-MTB are shown, including FIC predictions from INDIGO-MTB, Mtb-specific killing rate and interactions, number of cells/Mtb overtime used to evaluate simulated EBA, spatial analysis of antibiotic concentration and interactions, and sterilization time distributions from a collection of granulomas.

Next, to predict *in vivo* interactions and efficacy, here we integrate INDIGO-MTB predicted drug interactions within an existing multi-scale model of pathogen-immune dynamics leading to granuloma formation, known as *GranSim*^22,25–28^. *GranSim* integrates spatio-temporal host immunity, pathogen growth and drug PK/PD into a single computational framework. *GranSim* uses a hybrid agent-based model to describe interactions between immune cells and cytokines with bacteria and antibiotic delivery to granulomas, and provides a dynamic picture of pathogen clearance leading to granuloma sterilization^22^. Previously, we modeled PK/PD in *GranSim* and explored regimens with isoniazid and rifampin, three fluoroquinolones, and more recently HRZE^29–31^. However, in these studies we assumed no interaction between antibiotics. Hence, we now integrate INDIGO-MTB predictions of drug interactions within the *GranSim* framework, allowing the full characterization of drug interaction dynamics at the molecular and cellular scales, and verify our results against patient-level data. This allows us to evaluate how drug interactions and PK/PD at the molecular scale influence *in vivo* efficacy at the granuloma scale.

Our study herein represents the first pipeline that incorporates both *in vitro* drug interactions and *in vivo* PK/PD to simulate treatment dynamics of numerous drug regimens. Our study overcomes the limitation of prior studies that have focused on variation in PK/PD parameters alone to predict treatment outcome^32–34^. Combining INDIGO-MTB with *GranSim* allows us to compare different regimens based on the impact of their interactions on various simulated metrics such as rate of pathogen load decline in granulomas and granuloma sterilization rates. Our approach provides a measurement of drug interactions within lung granulomas based on concentrations that different bacterial populations are experiencing in their individual granuloma environment.

## Results

### Drug interactions significantly impact in vivo treatment dynamics in GranSim

We focus on combinations of 2, 3 or 4 drugs involving the first-line antibiotics and two fluoroquinolones (Supplementary Table 1). These drugs include isoniazid (H), rifampin (R), ethambutol (E), pyrazinamide (Z), moxifloxacin (M), and levofloxacin (L). We chose these drugs as they are part of the current standard-of-care for treating TB. Further, transcriptomics and PK/PD parameters are available for these drugs for simulation using both INDIGO-MTB and *GranSim*.

Using INDIGO-MTB, we first predicted all possible *in vitro* interaction outcomes for these combinations. The combinations are predicted to have FIC values that range from synergistic (e.g. HRZ – FIC of 0.74) to antagonistic (e.g. RM – FIC of 2.31). The standard regimen (HRZE) is predicted to be synergistic (FIC – 0.82) while moxifloxacin-containing regimens were mostly antagonistic (Supplementary Table 2).

Given the various factors that can impact antibiotic efficacy *in vivo* that are captured in *GranSim*, the relative impact of drug interactions on treatment outcomes is unclear. We hypothesized that analysis of various drug regimens with different drug interaction scores (FIC) can help tease out the impact on treatment outcome. Our previous studies of antibiotic treatment using *GranSim* did not consider drug interactions. Here we explore how either antagonistic or synergistic affects overall efficacy. We tested the impact of incorporating drug interactions on treatment dynamics using *GranSim*.

We input FIC values into *GranSim* and simulated the immune response and antibiotic delivery to granulomas (Methods). The plasma and tissue PK parameters for these drugs within the *GranSim* computational framework were derived from previous studies calibrating PK parameters to experimental plasma and lesion drug concentrations (Methods). For each regimen tested, 100 simulated granulomas were treated for up to 180 days with daily doses of each antibiotic in the specified regimen. To compare the efficacy of each of these regimens, we evaluate three measures: the log decrease in CFU per day, percent sterilization of granulomas, and average sterilization time.

The *in vitro* FIC value of each combination is correlated with each of the three simulated efficacy outcomes that we calculated (Figure 2). For our simulated log decrease in CFU per day from 0-14 days and the sterilization percent, we observe that both outcomes tend to decrease as FIC values go from synergistic to antagonistic (correlation R = −0.52 and −0.59 respectively, Figure 2A, 2B). The average sterilization time is positively correlated with FIC value (correlation R = 0.59, Figure 2). Overall, this indicates that synergistic regimens are more likely to sterilize a greater percentage of granulomas in a shorter time at both early and later time points.

**Figure 2.**
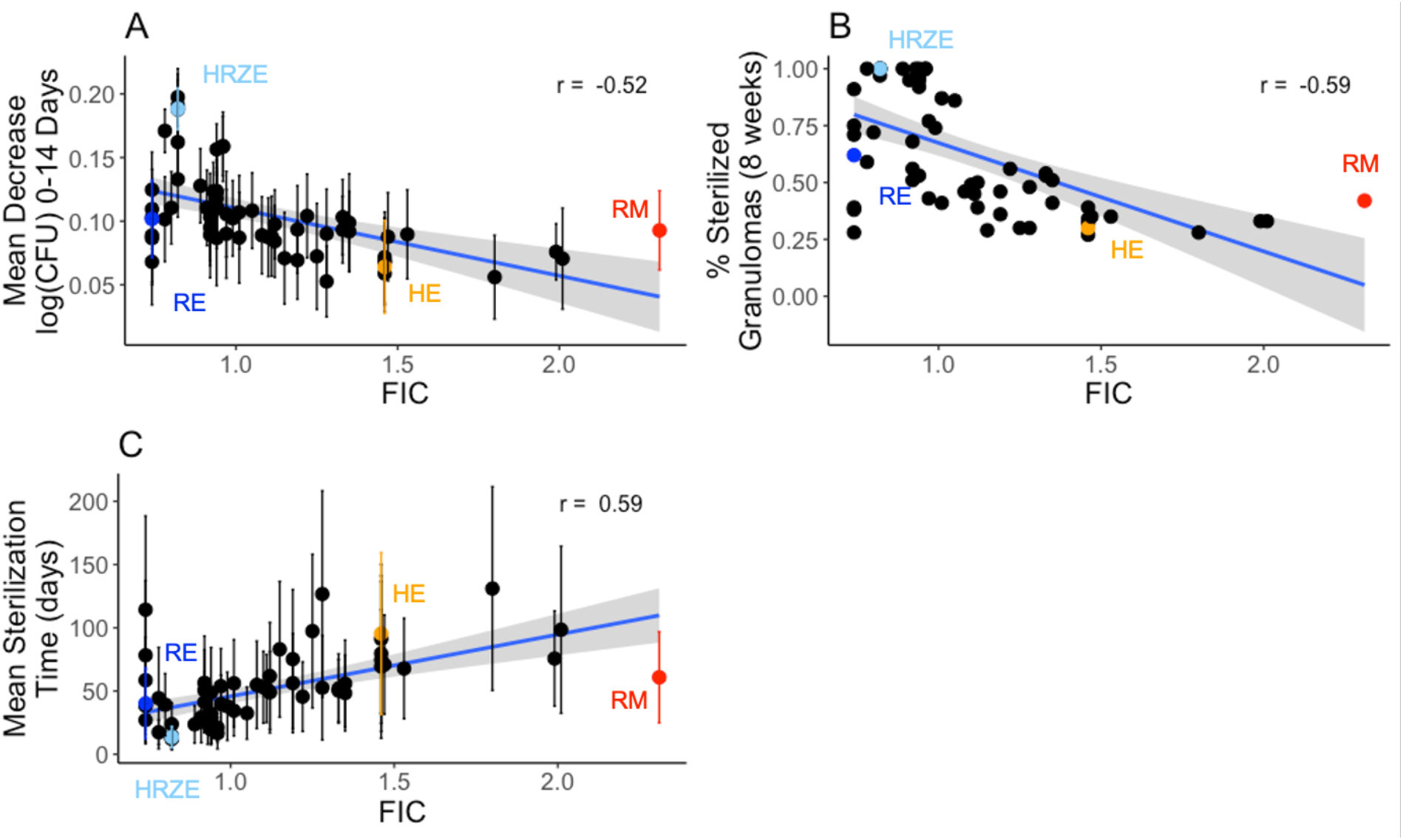
Regimen efficacy is correlated with FIC for 64 simulated drug regimens. Mean decrease in log CFU (0-14 days) averaged over 100 granulomas simulated for each drug regimen (A) and percentage of sterilized (negative) granulomas after eight weeks of treatment (B) are negatively correlated with FIC values, with correlation coefficients of −0.52 and −0.59, respectively. Mean sterilization time for each regimen over 100 granulomas (C) is positively correlated with FIC with a correlation coefficient of 0.59. Each point represents the regimen outcome measurement for a given regimen and error bars indicate +/- standard deviation from the sample of 100 granulomas simulated. The 64 drug regimens simulated are listed in Supplementary Table 1. The colored points correspond to the regimens HRZE (light blue), RE (dark blue), RM (red) and HE (orange) for emphasis.

Although these relationships show moderate levels of correlation, there are a few notable deviations. Interestingly, the regimen RM (FIC = 2.31) performs better than the less antagonistic regimen HE (FIC = 1.46). The best regimen in terms of average sterilization time is HRZE (FIC = 0.82); however, the regimen RE (FIC = 0.74) has a lower FIC but does not perform as well as HRZE. These results suggest that FIC values are not the only factor impacting granuloma sterilization. Because these results are based on sterilization in granulomas, the concentrations of each antibiotic in the granuloma compartment (based on dosage and PK) also impact the ability of each regimen to sterilize.

### The in vivo drug interaction score is predictive of treatment dynamics

The antibiotic killing rate is dependent not only on the FIC value, but also on local drug concentrations within a granuloma, the subpopulation of bacteria (intracellular, extracellular replicating, extracellular non-replicating), and the specific PD parameters of antibiotics involved. Based on the definition of the FIC, synergistic or antagonistic drug combinations result in a lower or higher effective concentration to achieve the same level of bacterial killing. To evaluate the overall impact of drug interactions on the calculated killing rate constant, we evaluate an *in vivo Drug Interaction Score* (iDIS) for the three subpopulations of bacteria. The iDIS measures the relative increase or decrease of the antibiotic killing rate constant due to the specific drug interaction. We calculate iDIS as the ratio of the antibiotic killing rate constant evaluated in the simulation to the rate constant if the interaction is simply additive (FIC equal to 1.0). This ratio provides a measure of how much the drug interaction impacts the killing rate constant and is unique for each individual mycobacterium within *GranSim* as drug concentrations change over time. At each time step during treatment, the average iDIS over all Mtb by subpopulation is evaluated as a model output.

Figure 3 shows the average iDIS for non-replicating Mtb over the first dose interval for each regimen, and its relationship to regimen outcomes. A value of 1 indicates the interaction has no impact on the killing rate constant; values greater than 1 or less than 1 indicate synergistic or antagonistic combinations, respectively.

**Figure 3.**
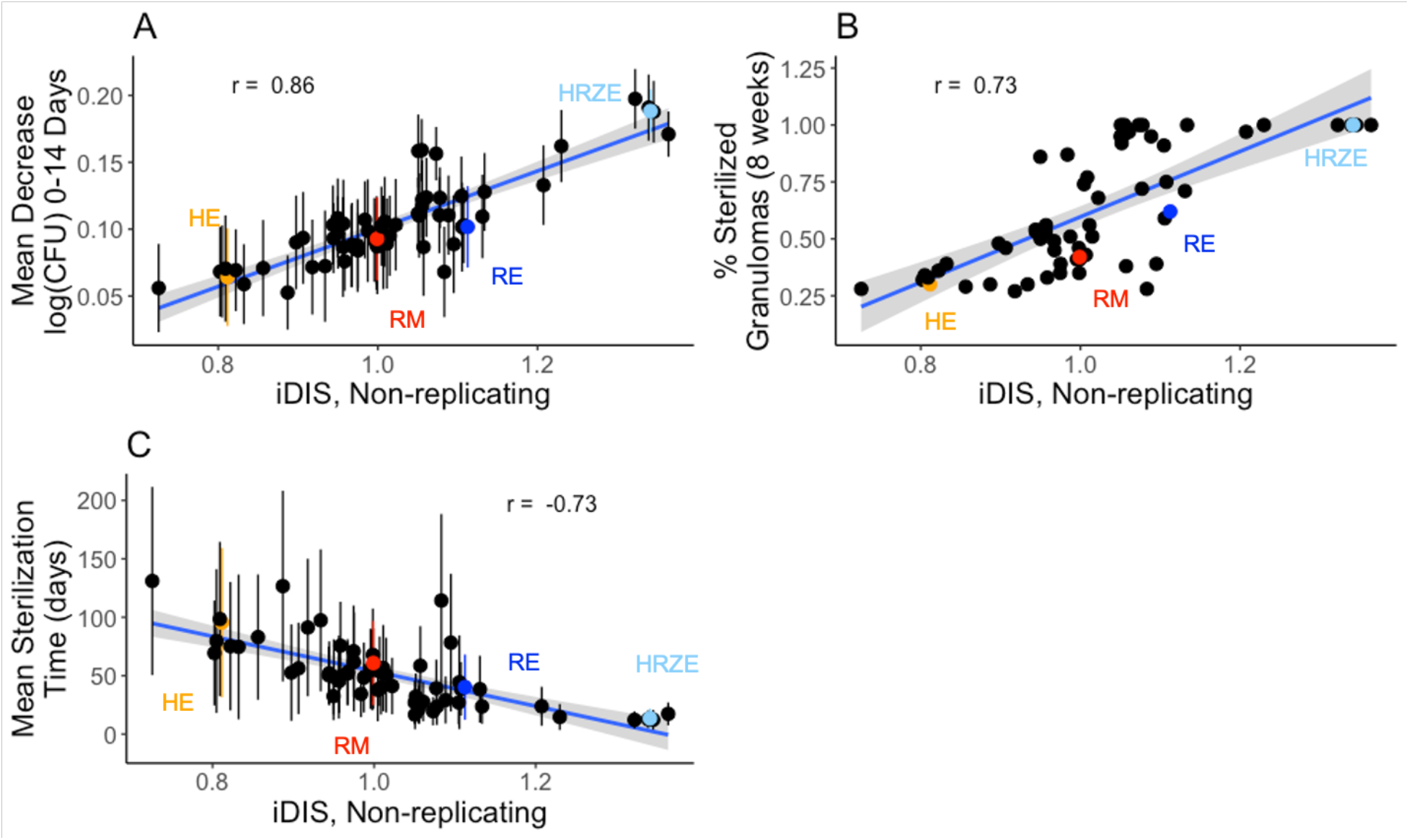
Regimen efficacy is correlated with the *in vivo* Drug Interaction Score (iDIS). iDIS associated with non-replicating Mtb killing is evaluated for 3 measures over 64 simulated drug combination regimens. The decrease in log CFU (0-14) averaged over 100 granulomas simulated for each regimen (A) and percentage of sterilized (negative) granulomas after eight weeks of treatment (B) are positively correlated with iDIS of non-replicating Mtb during the first 24 hours of treatment (correlation coefficients of 0.86 and 0.73, respectively). Mean sterilization time for each regimen over 100 granulomas (C) is negatively correlated with iDIS of non-replicating Mtb (correlation coefficient of −0.73). Each point represents the regimen outcome measurement for a given regimen and error bars indicate +/- standard deviation from the sample of 100 granulomas simulated. The 64 drug regimens simulated are listed in Supplementary Table 1. The colored points correspond to the regimens HRZE (light blue), RE (dark blue), RM (red) and HE (orange) for emphasis.

The iDIS for each regimen is strongly correlated with the outcomes from our *GranSim* simulations: log decrease in CFU per day (R = 0.86), percentage of negative granulomas at eight weeks (R = 0.73), and the average sterilization time (R = −0.73) (Figure 3). The correlations are much stronger than those observed for FIC (Figure 2), indicating that measuring the iDIS, which is calculated for specific granuloma environments, provides more information on regimen efficacy than examining FIC values, which are calculated based on *in vitro* environments.

Each combination of antibiotics has a different absolute killing rate constant based on the specific combination of PD parameters associated with that combination together with the distribution of antibiotics within a granuloma. These results suggest that iDIS provides a more accurate representation of how well a given combination of antibiotics achieves sterilization as it accounts for the unique killing rate constant that each individual Mtb experiences and measures the impact that an FIC value has on that killing rate constant.

Antibiotics work stronger on replicating Mtb than against non-replicating Mtb. Antibiotic killing rate constants that are higher and closer to their overall E_max_ value are less impacted by drug interactions. We found that correlations between regimen outcomes and the average iDIS associated with replicating extracellular and intracellular Mtb are weaker than when comparing regimen outcomes to the iDIS from non-replicating Mtb (Supplementary Figures 1 and 2). The average iDIS measurements for replicating Mtb are closer to 1.0 and weaken the correlation with regimen outcomes. Hence, the strong correlation between drug interactions and clinical outcomes are primarily driven by drug action against non-replicating bacteria.

Figure 4 shows a heat map of the mean sterilization time, iDIS and FIC for each regimen, ordered by decreasing iDIS. In general, the regimens with the fastest sterilization times also have high iDIS. The top 17 regimens, as measured by shortest average sterilization time, all contain RIF, indicating that RIF is a very important addition to regimens. Another general trend is that two-drug combinations typically perform worse than 3- or 4-drug combinations. Fluoroquinolones tend to participate in more antagonistic combinations, as measured by the iDIS. For example, 22 of the 31 MXF or LVX containing regimens are above the median iDIS of all 64 regimens. Two regimens (R23.5E45dpw2 and R23.5E90dpw1) with synergistic iDIS measurements showed slow sterilization times, as they were dosed less frequently than a day.

**Figure 4.**
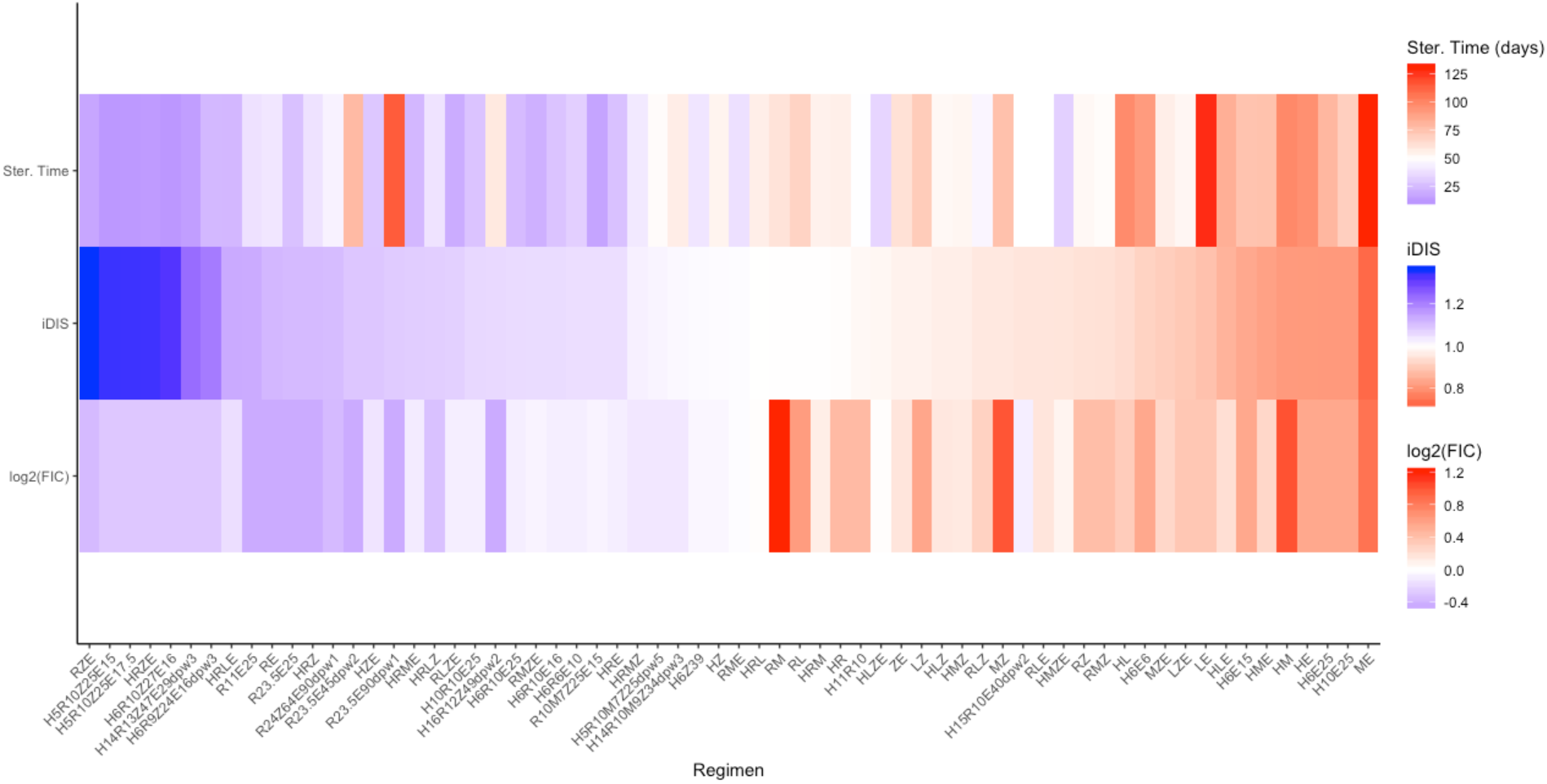
Heat map capturing three metrics for 64 different regimens. The list of regimens is ordered by decreasing predicted iDIS (middle row). For each regimen, the log2(FIC) value (bottom row) and the average predicted granuloma sterilization time (top row) are also represented. For predicted iDIS and FIC, blue represents synergy, white represents additivity, and red represents antagonism. For sterilization time, blue represents shorter sterilization times and red represents longer.

### INDIGO-MTB - GranSim regimen rankings are correlated with clinical rankings

To explore how our predictions of regimen efficacy compare to clinical results, we compare our INDIGO-MTB - *GranSim* treatment simulations to results from TB drug clinical trials. Drawing from the meta-analysis of phase II trials presented in Bonnet *et al* (2017), we selected all regimens that reported sputum culture conversion in solid media^35^. The efficacy metric presented for Phase IIb trials is the percent of patients with negative sputum culture after 8 weeks of therapy. Since our simulations predict treatment outcomes at the granuloma scale, we estimated how sterilization at the granuloma level relates to host-level culture conversion. Supplementary Figure 3 shows the comparison of the upper and lower bound estimates for percent sputum conversion from our simulation to clinical trial results for 26 regimens (Methods). For most regimens, estimates of sterilization percentage compare closely to clinically measured culture conversion. In general, *INDIGO-MTB - GranSim* simulations appear to overpredict the rates of sterilization and most incorrect predictions fall into this category (Supplementary Figure 3). This observed overprediction is likely due to the simplification of predicting sterilization at the granuloma scale that does not include the full spectrum of complex granuloma lesions, failed adherence to regimens, and other factors that complicate TB treatment.

We next validate our simulation results by comparing the ranking of the efficacy of each of the regimens with the corresponding ranking of the efficacy from clinical trials (Figure 5). The clinical rank is determined by ranking each regimen by the pooled culture conversion after 8 weeks, so that a ranking of 1 is the regimen with the lowest culture conversion. The simulation rank is determined by percentage of granulomas sterilized after 8 weeks. We used the Spearman ranked correlation coefficient, weighted by the number of patients in each pooled regimen result, and found a significantly strong correlation between simulations and clinical trials (R = 0.72).

**Figure 5.**
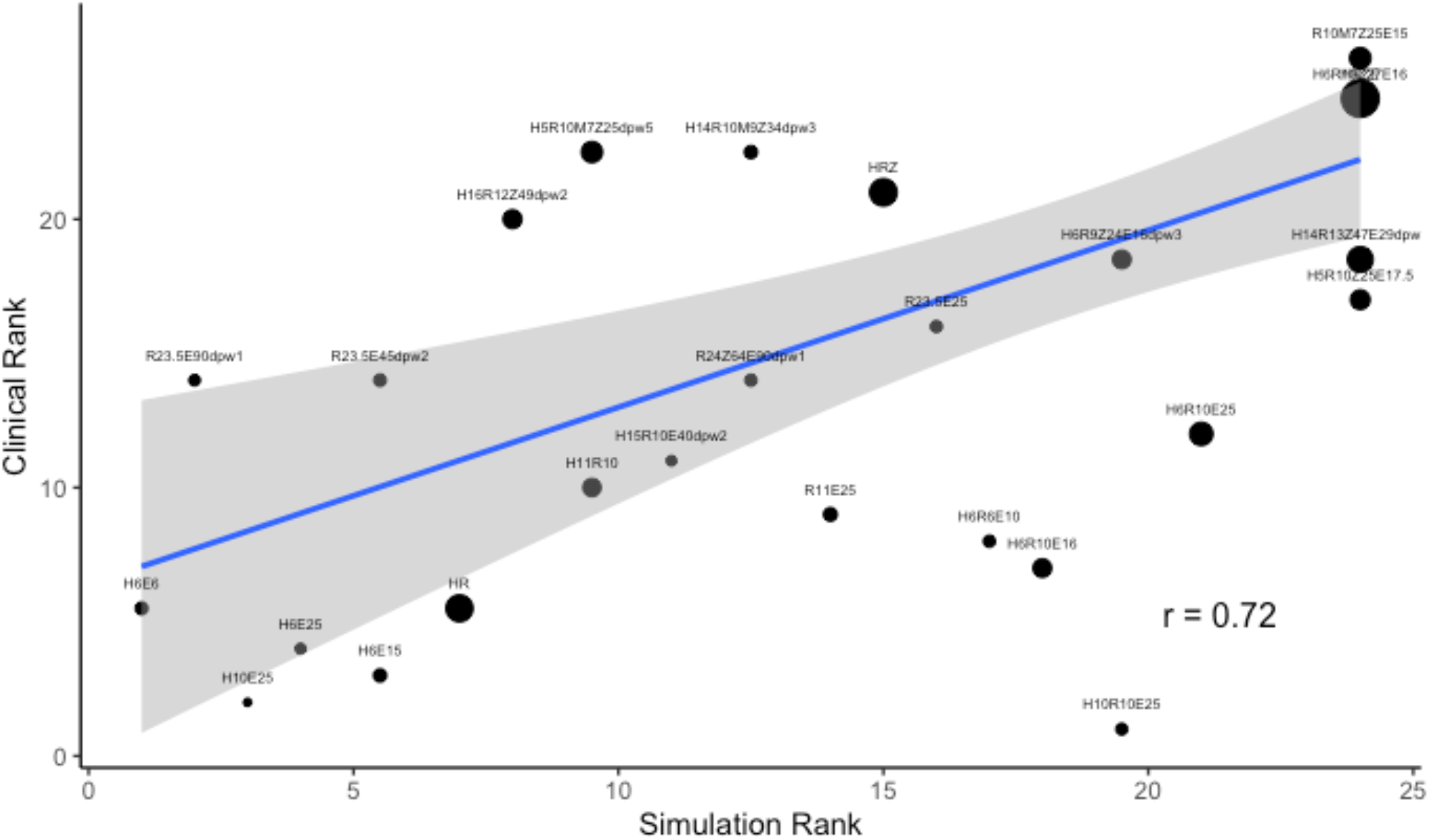
INDIGO-MTB-GranSim compared with clinical data. Comparison and validation of treatment simulations with clinical trial results for 26 different regimens compiled in Bonnet *et al*. (2017)^35^. Predictions from *GranSim* simulations for 26 drug regimens correlate with clinical outcomes. The simulation rank, ranked by percentage of sterilized granulomas after 8 weeks, and clinical rank, ranked by clinically reported culture conversion, have a weighted correlation of 0.72, weighted by the number of patients treated with each regimen.

Recent phase III clinical trials have investigated the impact of introducing fluoroquinolones into treatment regimens to treat drug-susceptible TB, some with the additional intent of shortening treatment time from six months to four. Many of these trials have failed to show improvement in TB treatment, and often led to higher rates of unfavorable outcomes at the trial’s endpoints^36–38^. These trends are reflected in our analysis of the drug interactions for various drug combinations. The control regimen, HRZE, is strongly synergistic as measured by iDIS, and we predict short average sterilization times (14 days, Figure 4). In contrast, fluoroquinolone containing regimens, such as HRMZ and RMZE, are closer to additive, and are predicted to have longer sterilization times of 41 and 21 days respectively. These trends indicate that our simulation predictions are consistent with phase III clinical trial observations. Thus INDIGO-MTB - *GranSim* simulations provide strong predictive measures of clinical outcome for different regimens.

Both iDIS and INDIGO-MTB FIC predictions for these regimens are also significantly correlated with their clinical ranking (Supplementary Figure 3B and 3C). These simulated results are consistent with the correlation observed in a prior study between INDIGO-MTB FIC scores and percentage of patients with negative culture after treatment from 57 randomized clinical trials^8^. Based on these results, both interaction measurements have the potential to help predict the clinical efficacy of drug combination regimens.

The FIC value is one measure of drug interactions *in vitro*; however, there are many factors that impact regimen efficacy *in vivo* that FIC alone does not capture. Measuring the iDIS can incorporate changes in concentration due to PK variability, changes to dosing regimens, and heterogeneous antibiotic concentrations due to granuloma structure and the varying environments where bacteria reside impact the degree to which the drug interactions impact killing rates. It also includes effects of the immune responses occurring with granulomas.

### Spatial variation in drug concentration influences iDIS in granulomas

Nonuniform drug distributions within granulomas arise due to barriers to diffusion that the cellular structure of granulomas creates^20–22,29^. The spatial variation in antibiotic concentrations within a granuloma leads to variations in local effective concentrations, and ultimately antibiotic killing rates and iDIS. The free drug concentrations available to induce bactericidal activity against Mtb are also influenced by binding to extracellular matrix as well as partitioning into macrophages^39^. Hence, we next focused on the contribution of the drug spatial variation to iDIS.

Figure 6 shows the spatial variation for effective drug concentrations normalized to the regimen’s non-replicating Mtb *C*_50_ parameter and iDIS for four of the regimens simulated: HRZE, RE, HE and RM. These regimens demonstrate a mix of synergistic and antagonistic combinations that also exhibit both strong and weak iDIS values. For the two synergistic combinations, HRZE and RE, effective concentrations are lower within a granuloma than just outside it, lowering the killing rate constants. Because most non-replicating Mtb are found within the hypoxic caseum which is typically located toward the center of the granuloma, the effective concentrations in those caseated regions and inside the granuloma are the concentrations that are relevant to predicting sterilization rates. However, lower effective concentrations inside granulomas and caseated regions can also result in higher iDIS (Figure 6). Because iDIS tends to be higher within granulomas where Mtb reside, this may contribute to synergistic combinations performing better than antagonistic combinations of similar effective concentrations.

**Figure 6.**
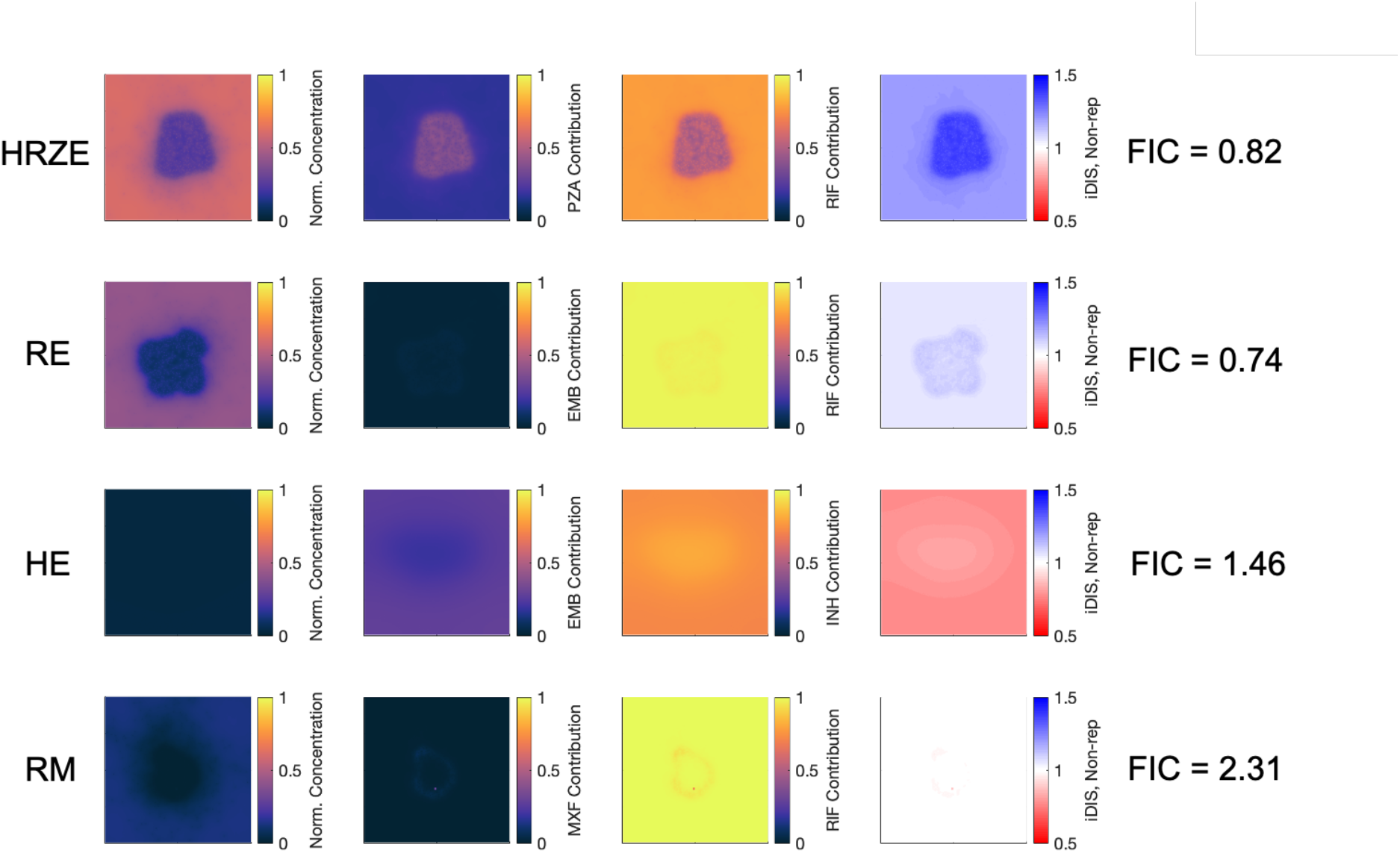
Contribution of individual antibiotics to the *in vivo* drug interaction score (iDIS). Values of iDIS are associated with proportion of each antibiotic’s contribution to the effective concentration. Four different regimens are shown: HRZE (first row), RE (second row), HE (third row), and RM (fourth row). Heat maps show the effective concentration normalized to the C_50_ for non-replicating Mtb of the combination (first column) and the fraction of each antibiotic’s contribution to the effective concentration (second and third columns). The calculated iDIS value for non-replicating Mtb is shown in the fourth column, with the color bar representing the iDIS value with blue representing a synergy, white representing additivity, and red representing antagonism. All heat maps reflect conditions 6 hours after dosage with each antibiotic in the relevant regimen.

An additional aspect that influences the iDIS is the relative contribution of each antibiotic in the combinations (Figure 6, columns 2 and 3). Antibiotic combinations that contribute more equally to the effective concentration deviate more from additivity than combinations of antibiotic concentrations in which contributions are uneven (i.e. when one antibiotic has a much higher adjusted concentration than the other antibiotics). Although RE has a more synergistic FIC than HRZE, the iDIS against non-replicating Mtb for HRZE is higher than RE, and HRZE has better efficacy than RE. This is partially due to R and Z contributing to the effective concentration in the granuloma evenly (R ~50% contribution, Z ~40% contribution). The contributions from R and E in RE, however, are more disproportionate, with >90% of the effective concentration due to R and <10% of the effective concentration from E.

A similar situation occurs with the two antagonistic combinations HE and RM. Although RM is predicted to be strongly antagonistic, its efficacy is still average compared across all regimens tested. HE, on the other hand, has a less antagonistic FIC, but performs more poorly than RM. When looking at both antibiotic contributions for these two regimens and the iDIS against non-replicating Mtb, we see that H accounts for ~75% of the effective concentration and E accounts for ~25%. Although not equal contributions, this still allows for an antagonistic interaction to occur. With the RM combination, R accounts for almost all of the contribution to the effective concentration because its levels are higher relative to its own *C*_50_, resulting in almost no antagonistic interaction to occur with M, despite the high FIC. For antagonistic combinations, uneven contributions from the different antibiotics in the combination can mitigate the effect of the antagonistic interaction.

Due to this dependence on drug concentrations, the predicted iDIS varies for different doses and regimens of the same drug combination (Supplementary Figure 4). In contrast, the FIC interaction score is fixed for a combination irrespective of the dosage. As doses vary *in vivo*, the strength of the synergistic or antagonistic interactions can either increase or decrease, depending on the specific combination of antibiotics. One common trend is less frequent dosing tends to decrease the interaction strength, which we observe for both the HRZE and RE combinations.

### Plasma clearance rates correlate with in vivo drug interaction score

Even for a given simulated granuloma and drug regimen, antibiotic concentrations can vary due to host-to-host PK variability ^31^. This can also result in changes in iDIS. We picked four different regimens, ranging from synergistic to antagonistic (HRZE, RE, HE, RM) to exhaustively explore the impact of various PK parameters. The plasma clearance rate constant for many of the antibiotics in these regimens is significantly correlated with predicted iDIS, particularly the clearance rate constants for R, E and M (Table 1). A more detailed analysis of correlations between plasma PK parameters and iDIS is shown in Supplementary Table 3. The R clearance rate constant is strongly correlated with iDIS, with coefficients between 0.8 and 0.9, depending on the regimen. The correlation coefficients for the clearance rate constant for R are positive for synergistic combinations (HRZE, RE), and negative for antagonistic combinations (RM). Because iDIS values that deviate further from 1 imply a stronger interaction, this means that faster clearance rates for R tend to increase the interaction strength. The opposite is true for the clearance rate constant of E. The correlation coefficient between the clearance rate constant for E and predicted iDIS is negative in synergistic combinations (HRZE, RE), and positive in the antagonistic combination (HE). While this may seem counterintuitive, it supports the idea that the iDIS value is dependent on relative in vivo drug concentrations. Faster clearance rates generally result in lower concentrations in plasma, and consequently lower concentrations in the granuloma. Because R tends to contribute more to effective concentrations than other antibiotics, increasing the clearance rates for R will strengthen the interaction by decreasing R concentration and allow for more even contributions. For E, whose contribution to effective concentration tends to be lower, increasing clearance rates result in lower E concentrations and contributions, and scenarios of even more lopsided contributions and less interaction. Relative drug concentrations inside the granuloma affect the strength of drug interactions, and these strong correlations indicate that interactions may be stronger or weaker for certain combinations depending on an individual’s PK^31^.

**Table 1:** Significant antibiotic clearance rate constants in determining iDIS. The relationship between clearance rate constants for different antibiotics is correlated with iDIS with non-replicating Mtb during the first dose of therapy. Table shows PRCC values relating the clearance rate constants to the predicted iDIS for the regimens HRZE, RE, HE and RM. The values shown represent PRCC values that are significant with p < 0.01.

| Regimen | PRCC Values for predicted iDIS |  |  |
| --- | --- | --- | --- |
|  | R clearance | E clearance | M clearance |
| HRZE | 0.89 | -0.56 | N/A |
| RE | 0.90 | -0.92 | N/A |
| HE | N/A | 0.88 | N/A |
| RM | -0.83 | N/A | 0.99 |

## Discussion

The need for multiple drug regimens to treat TB raises a number of issues that can be capitalized on to advance treatment for the world’s worst killer by disease. In particular, understanding the role of interactions occurring in either a synergistic or antagonistic fashion between anti-TB antibiotics *in vivo* may provide a more rational approach to choosing novel combinations that have greater clinical efficacy. However, measuring drug interactions *in vivo* is challenging due to the limited throughput, cost, and time involved in testing drugs in model organisms. In this study, we introduce a computational pipeline that can simulate *in vivo* interactions by using datasets derived from individual drug-response transcriptomes and PK/PD, thereby greatly reducing cost and time. Our approach integrates interaction scores of combinations of antibiotics into a computational model that simulates drug delivery into the lung, spatial concentrations of drugs and pharmacodynamic effects within TB granulomas.

To evaluate drug regimens, we introduce a new metric called the *in vivo* drug interaction score (iDIS) that is dynamic and unique for each mycobacterium based on its location and metabolic state (i.e. replicating/non-replicating) within a granuloma. Unlike the *in vitro* drug interaction scores derived from checkerboard assays and INDIGO-MTB, which are fixed for given drug combinations, the *in vivo* score can provide a more nuanced impact of drug interactions on pathogen clearance. This allowed us to compare various drug regimens and rank them based on their *in vivo* interactions. We found that our ranking of regimens is highly concordant with clinically observed efficacy of various drug combinations^35^.

When simulating all regimens considered in this study, the FIC alone does correlate with the simulated and predicted outcomes from our model. However, there are a handful of regimens that do not fit the trend. Measuring the iDIS, which evaluates the relative increase or decrease in killing rate due to the interaction within the complex granuloma environment, provides a complementary measure of regimen outcome. This is because the effect that drug interactions have on killing rate constants is dependent on the balance of contributions from each single antibiotic. Combinations with highly synergistic or antagonistic FIC values may be closer to additive if only one antibiotic is present in sufficient quantities within a granuloma.

Our analysis of various drug regimens revealed ways of amplifying synergy as well mechanisms to mitigate antagonism *in vivo*. We found that some combinations with *in vitro* antagonism perform well clinically due to the distinct spatial distribution of the underlying drugs. Antagonistic interactions can be mitigated if the drugs have uneven distributions and effective concentrations or through less frequent dosing. Overall, we find that combinations with strong *in vitro* synergy remain synergistic or additive *in vivo*. Hence screening for synergy *in vitro* can be a useful strategy for identifying regimens with strong *in vivo* activity. In a minority of cases, this synergy may not be achieved *in vivo*; nevertheless, synergistic combinations generally outperform antagonistic regimens.

As part of this study, we wanted to determine which combinations of antibiotics are predicted to have strong synergy and antagonism, as well as which combinations are predicted to have high efficacy. We screened 64 combinations and regimens of front-line regimens (HRZE) along with M and L. The clinically used HRZE regimen does outperform other screened combinations, which highlights the need for new drugs to achieve the aim of improving TB treatment. Based on the INDIGO-MTB model, we previously identified drug combinations involving new TB drugs such as bedaquiline that have better synergy than HRZE. Given the concordance between INDIGO-MTB FIC and various clinical metrics observed in this study, the synergistic combinations identified by INDIGO-MTB may be promising leads for further optimization using *GranSim* for reducing treatment time^16^.

A limitation to our computational model is that the same FIC value is applied to all Mtb within a simulation, regardless of its environment or metabolic state. It is likely that the strength of a given interaction, or even whether it is synergistic or antagonistic, is dependent on the bacterium’s microenvironment in the granuloma^40,41^. Additionally, these simulations represent treatment of primary granulomas in TB disease, and do not necessarily reflect the true and enormous complexity of granuloma lesions that occur during TB disease. Further, as the simulations are at the granuloma scale, relating the outcomes measured by the simulation to clinical outcomes is difficult. The final limitation is that we only considered combinations of six different antibiotics. There are many other antibiotics in use or in development for use to treat TB. Expanding our ability to accurately simulate the PK/PD of additional antibiotics will greatly increase our ability to answer how important drug interactions are in determining regimen efficacy.

In sum, our study addresses an important gap in current methods for identifying promising drug combinations for TB treatment by presenting a new pipeline for evaluating interactions between drugs *in vivo*. This pipeline provides an additional metric with which to evaluate novel combinations of antibiotics, explain mechanisms of failed regimens, and assist in optimization regimens as we expand our list of potential antibiotics.

## Methods

### INDIGO-MTB model for predicting drug interactions

INDIGO-MTB identifies interactions between drugs in a combination regimen by utilizing pathogen transcriptomics in response to individual drugs. INDIGO-MTB was built using drug response transcriptome data for 164 drugs, including well known drugs rifampicin, isoniazid, streptomycin, and several fluoroquinolones^8^. The model first generates a drug-gene association network using the transcriptomics data, and the machine-learning algorithm, Random Forest. The algorithm identifies genes that are predictive of drug interaction outcomes using a training data set of known interactions^42^. This trained network model is used to predict interactions for novel drug combinations and provides the Fractional Inhibitory Concentration (FIC) as an output (Figure 1). The model can identify all possible 2-way, 3-way, 4-way and 5-way synergistic, additive and antagonistic drug interactions after *in silico* screening of more than 1 million potential drug combinations. INDIGO-MTB predicted FIC scores to be integrated within *GranSim* were generated for all possible combinations of the first line drugs and two fluoroquinolones (H, R, Z, E, Levofloxacin (L) and Moxifloxacin (M)), as listed in Supplementary Table 1.

### GranSim model of granuloma formation and function

*GranSim* is a well-established agent-based model of granuloma formation and function^22,25–28^. It simulates the spatial heterogeneity and bacterial burden of primary TB lesions by simulating the immune response to infection with Mtb in a computational grid representing a small section of lung tissue, with formation of a granuloma as an emergent behavior (Figure 1). The simulation begins with a single infected macrophage at the center of the grid, and macrophages and T cells are recruited to the site of infection and interact with each other according to immunology-based rules that describe cell movement, activation, cytokine secretion, and killing of bacteria (for a full list of rules see referenced webpage^43^). Bacteria are tracked individually and modeled as individual agents in the simulation, existing in three distinct subpopulations: intracellular (inside macrophages), extracellular replicating and extracellular non-replicating. The effective growth rates of extracellular bacteria are modulated by the number of bacteria in a given grid compartment. The growth rate becomes zero when the carrying capacity for that compartment is reached to reflect the relative availability of nutrients and physical space limitations^27^. Growth rates of extracellular Mtb are also slowed by the presence of caseum (dead cell debris), as a way to estimate the effect of lack of oxygen^44^. The parameter values describing rules and interactions are based on previous *GranSim* studies and evidence from experimental literature and datasets on non-human primates^22,31,45^.

### Simulation of antibiotic delivery and concentrations within granuloma

We simulate antibiotic delivery within the *GranSim* computational framework as previously described^22,29,31^. Briefly, a plasma PK model simulates absorption into plasma following an oral dose, exchange with peripheral tissue, and first-order elimination from plasma. Flux of antibiotics into the simulation grid is based on the local gradient between the average drug concentration surrounding vascular sources on the agent-based grid and the plasma concentration. Here we use a 200×200 grid representing a 4 mm x 4 mm lung section. The flux is calculated over time as plasma concentrations change within and around each vascular source and allows for delivery or subtraction from the computational lung environment, depending on the direction of the concentration gradient. Once on the grid, antibiotics diffuse, bind to extracellular material (epithelial tissue and caseum), partition within macrophages and degrade (Figure 1(b)). Based on relative binding and partitioning rates into macrophages, concentrations of intracellular and bound antibiotic are modeled at pseudo-steady state for isoniazid, rifampin, ethambutol and pyrazinamide. The drugs moxifloxacin and levofloxacin exhibit slower rates of binding and partitioning relative to diffusion. Hence the dynamic binding and partitioning of these drugs are modeled using ordinary differential equations^29^. We determined plasma PK parameters by calibration to human data as previously described^20,31,46^. We calibrated tissue PK parameters based on concentrations in rabbit or human lesions^20,29,31,39^.

### Calculation of antibiotic killing rate and in vivo drug interaction

We calculate the antibiotic killing rate constant using an Emax model (Hill equation) as we have done previously^22^. This antibiotic killing rate constant is evaluated at each time step for every Mtb in the simulation based on the local grid concentrations as they change over time. The antibiotic killing rate constant (*k*) is evaluated as

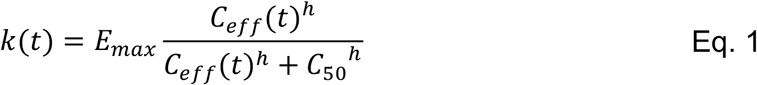

where *E_max_* is the maximal killing rate constant, *C*_50_ is the concentration at which half maximal killing is achieved, *h* is the Hill coefficient, and *C_eff_* is the effective concentration of the antibiotic (or combination of antibiotics). To reflect each antibiotic’s unique levels of activity against different sub-populations of bacteria, the PD parameters *C*_50_, *E_max_* and *h* vary depending on the location of the bacteria within the granuloma (intracellular, extracellular replicating, or extracellular non-replicating).The relationship between a combination of drug concentrations and pharmacodynamic effect (such as killing and inhibition) is described using the Loewe Additivity model^14,47^. In the Loewe additivity model, a simply additive interaction between two antibiotics is described by

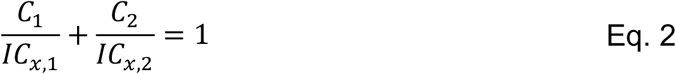

where *IC*_*x*,1_ and *IC*_*x*,2_ are the inhibitory concentrations of drugs 1 and 2 that achieve x% inhibition on their own, and *C*_1_ and *C*_2_ are the concentrations that achieve the same level of inhibition in combination. We can convert the concentration of drug 2 to an equipotent concentration of drug 1, shown in Figure 7 and denoted *C*_2,*adj*_. This gives the concentration of drug 1 that results in the same antibiotic killing rate constant as the given concentration of drug 2 (*C*_2_), which we define as the adjusted concentration for drug 2 (*C*_2,*adj*_).

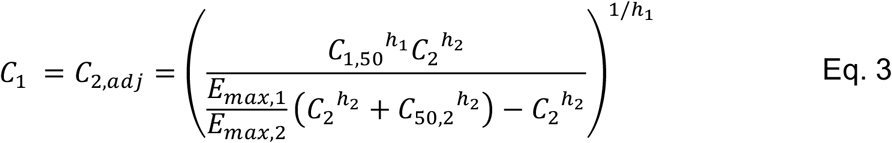

**Figure 7:**
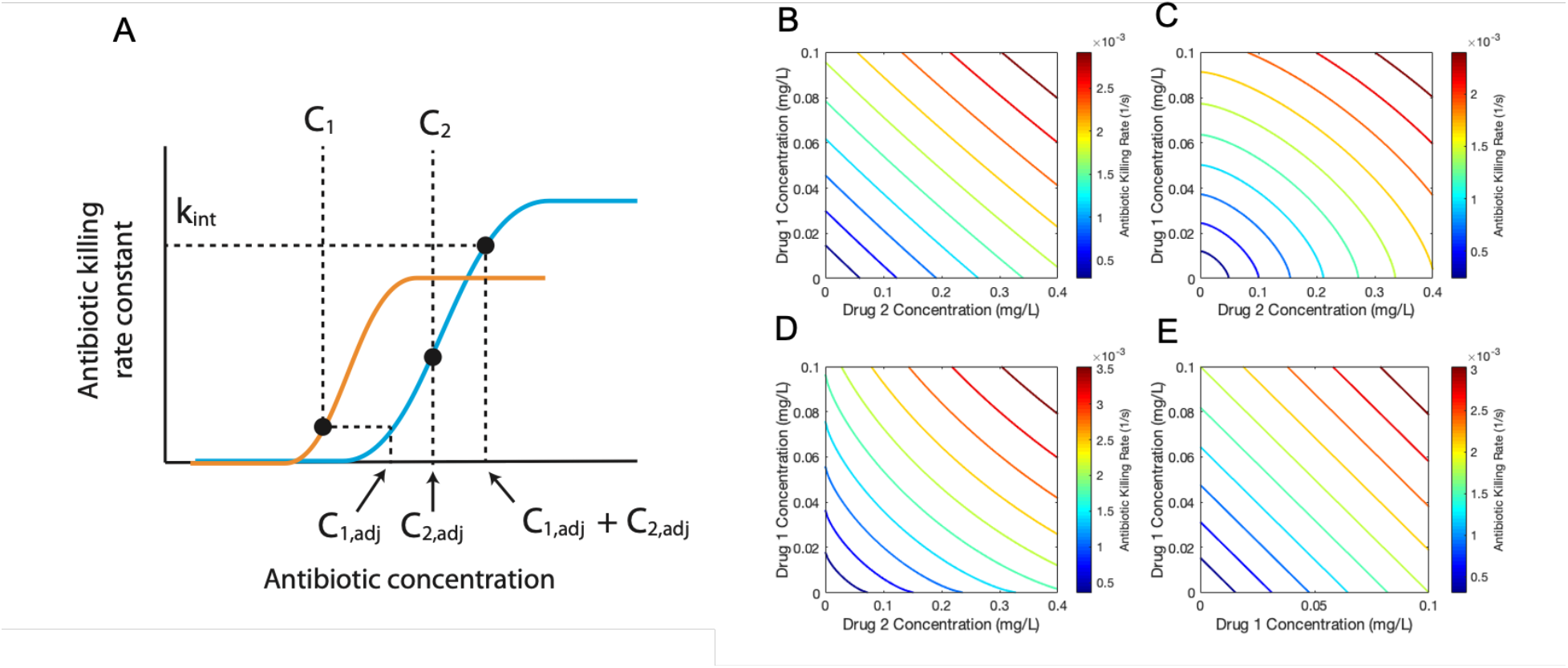
Graphical representation of computing the adjusted concentration and killing rate constant. (Eq. 1 and 3). The adjusted concentration of a drug is found by computing the equipotent concentration for another drug. The plot of two Hill curves for three different drugs (drug 1, orange; drug 2, blue) shows the relationship between concentrations of the two antibiotics and their adjusted concentration (A). The effective concentration, evaluated as the sum of the adjusted concentrations, determines the antibiotic killing rate constant. Antibiotic killing rate constant contours show the behavior of the drug interaction model for a combination of two theoretical drugs. Drug 1 has a c50 of 1 mg/L, Emax of 0.02 1/s, and a hill coefficient of 1. Drug 2 has a c50 of 2 mg/L, Emax of 0.01 1/s, and a hill coefficient of 1. When the two drugs have an FIC of 1.0 (B), 1.5 (C), or 0.6 (D), the contours show the characteristic straight line or curved contours characteristic of checkerboard assays for additive, antagonistic, or synergistic combinations. A sham combination of Drug 1 (E) results in a simply additive case.

The corresponding inhibitory concentrations for a given x% inhibition for drugs 1 and 2 are now both equivalent to *IC*_*x*,1_, because both *C*_1_ and *C*_2,*adj*_ are expressed in terms of concentration of drug 1. Substituting *C*_2,*adj*_ for *C*_2_ and *IC*_*x*,2_ for *IC*_*x*,1_, Equation Eq. 2 can be rewritten as

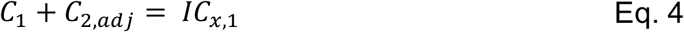

If there are 3 or more drugs under consideration, we define this sum of concentrations as the effective concentration (*C_eff_*) of a combination of *n* antibiotics:

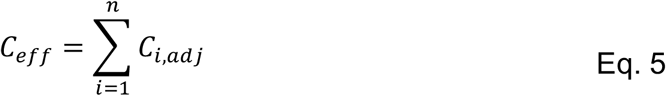

We define synergy or antagonism between two or more drugs based on deviations from simple additivity, as assumed above. This deviation is represented using the Fractional Inhibitory Concentration (FIC)^48^:

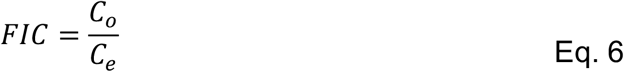

where *C_o_* represents the observed combined drug concentration to yield a given level of inhibition, and *C_e_* is the expected combined drug concentration to yield the same level of inhibition if the two drugs or more drugs were simply additive^14^. The FIC measures changes in potency, i.e. how much drug is needed to produce a certain pharmacodynamic effect. Based on the value of FIC, synergistic or antagonistic combinations result in a lower or higher effective drug concentration to achieve the same level of killing. To incorporate drug interactions into our model, we assume the effective concentration for a combination of *n* drugs is adjusted from Eq.5 based on the FIC value:

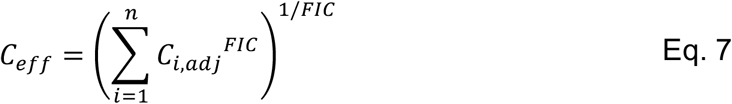

Eq. 7 adjusts effective concentration so that synergistic combinations (FIC < 1) result in a higher effective concentration, and antagonistic combinations (FIC > 1) result in a lower effective concentration. Using our defined effective concentration, we substitute Eq. 7 into Eq. 1 to evaluate the antibiotic killing rate constant for combinations of antibiotics while also accounting for drug interactions. Our drug interaction model and effective concentration formulae accurately recreate *in vitro* drug interaction behavior observed in checkerboard assays (Figure 7)^14^.

To evaluate the impact that each drug interaction has on the calculated killing rate constant (Eq. 1) for a given combination of antibiotics in our *in vivo* simulation, we define an *in vivo Drug Interaction Score* (iDIS). The iDIS is the ratio of the bacterial killing rate constant with a predicted FIC to the killing rate constant if FIC was equal to 1, i.e. no or additive drug interactions. This allows us to quantify the impact that drug interactions have on bacterial killing for each individual Mtb at each time step during simulated treatment.

### Antibiotic treatment simulations and calculation of regimen efficacy

To simulate treatment with different antibiotic combinations, we first created an *in silico* granuloma library to generate a set of granulomas. Each library consists of 500 granulomas simulations, generated from 100 parameter sets sampled with Latin Hypercube Sampling (LHS), and each parameter set was replicated five times^49,50^. Supplementary Table 2 lists the parameters varied and their ranges, which have been established in previous work^22,31^. Parameter ranges capture natural variability in the immune response and lung environment, such as differences in cellular recruitment and immune cell activation. In addition, replicating simulations with the same parameter set incorporates variability due to stochasticity in the simulations. Granulomas are simulated for 300 days in the absence of antibiotics. At day 300, a random sample of 100 unsterilized granulomas is selected from the relevant library for treatment. The prescribed regimens are simulated for 180 days or until the granuloma is sterilized. See Supplementary Table 1 for the full list of regimens tested.

We evaluate three metrics from our simulations to assess the efficacy of each regimen tested: log decrease in CFU per day, percent of simulated granulomas that are sterilized after eight weeks of treatment (sterilization percent), and average time at which those granulomas become sterile (sterilization time). For each regimen, 100 granulomas are simulated and results from those simulations are used to calculate 3 outcomes measures: simulated log decrease in CFU per day, sterilization percent, and sterilization time.

### Comparison to clinical trials

To validate our model results, we compared our treatment simulation outcomes to Phase IIb clinical trial data^35^. We compared the clinical datasets outcomes for each regimen with our simulated granuloma sterilizations after 8 weeks of treatment. We used the percent of granulomas that are completely sterilized at 8 weeks as a lower bound estimate. Our upper bound estimate is the percentage of granulomas with fewer than 10 CFU after 8 weeks. We chose this value as these granulomas with low bacterial load would not be detectable in sputum. Additionally, we compared the rank of clinically tested regimens, ranked by sputum conversion, to the rank of regimens based on simulation results. Simulated regimen rankings were ranked by average sterilization time, FIC, and average iDIS for non-replicating Mtb. Further, we compared our predicted treatment sterilization times for fluoroquinolone-containing regimens with clinical endpoints (up to 6 months) from recent phase III clinical trials that include fluoroquinolones for treating drug-susceptible TB^36–38^.

### Plasma PK sensitivity analysis on interaction strength

We performed a sensitivity analysis to evaluate how PK parameters impact the predicted iDIS for regimens with different levels of synergy. For four regimens (HRZE, RE, HE, RM), we selected a single granuloma to simulate treatment for one day to measure an iDIS. For each regimen, we simulated the granuloma 500 times with different plasma PK parameters sampled using LHS. For each plasma PK parameter set, we calculated the average iDIS over the first day of dosing over all non-replicating Mtb. Finally, we evaluated the partial ranked correlation coefficient (PRCC) between each plasma PK parameter and the predicted iDIS to determine the impact each parameter has on the drug interactions^50,51^.

## Acknowledgements

This research was supported by the following grants from the National Institutes of Health: U01HL131072 (to DEK, JJL) and R01AI123093 and R01AI150684 (awarded to DEK) and UL1TR002240, U19AI106761 (to SC), We also acknowledge funds from University of Michigan (UM) Precision Health (SC, DEK, JJL), UM Office of the Vice Provost of Research (SC) and UM MCUBED (SC, DEK). Simulations used resources of the National Energy Research Scientific Computing Center, which is supported by the Office of Science of the U.S. Department of Energy under Contract No. ACI-1053575 and the Extreme Science and Engineering Discovery Environment (XSEDE), which is supported by National Science Foundation grant MCB140228. We thank Paul Wolberg for computational assistance and support.

## Author Contributions

The simulations presented in this work were completed by JC. SC and AS provided FIC predictions from INDIGO-MTB. All authors (JC, AS, DE, JL, SC) contributed to the conceptualization of this work, analysis of results, and writing and editing.

## References

1. Dheda, K., Barry, C. E. & Maartens, G. Tuberculosis. Lancet 387, 1211–1226 (2016).

2. Global tuberculosis report 2019. (2019).

3. Zumla, A. et al. Tuberculosis treatment and management-an update on treatment regimens, trials, new drugs, and adjunct therapies. Lancet Respir. Med. 3, 220–234 (2015).

4. Mdluli, K., Kaneko, T. & Upton, A. The Tuberculosis Drug Discovery and Development Pipeline and Emerging Drug Targets. Cold Spring Harb Perspect Med 5, a021154 (2015).

5. Falzon, D. et al. World Health Organization treatment guidelines for drug-resistant tuberculosis, 2016 update. Eur. Respir. J. 49, (2017).

6. Conradie, F. et al. Treatment of highly drug-resistant pulmonary tuberculosis. N. Engl. J. Med. 382, 893–902 (2020).

7. Xu, J. et al. Contribution of pretomanid to novel regimens containing bedaquiline with either linezolid or moxifloxacin and pyrazinamide in murine models of tuberculosis. Antimicrob. Agents Chemother. 63, 1–14 (2019).

8. Ma, S. et al. Transcriptomic signatures predict regulators of drug synergy and clinical regimen efficacy against tuberculosis. MBio 10, 1–16 (2019).

9. Berenbaum, M. C. A method for testing for synergy with any number of agents. J. Infect. Dis. 137, 122–130 (1978).

10. Berenbaum, M. C. What is Synergy? Pharmacol. Rev. 1989, 93–141 (1989).

11. Michel, J. B., Yeh, P. J., Chait, R., Moellering, R. C. & Kishony, R. Drug interactions modulate the potential for evolution of resistance. PNAS 105, 14918–14923 (2008).

12. Chandrasekaran, S. et al. Chemogenomics and orthology-based design of antibiotic combination therapies. Mol. Syst. Biol. 12, 872 (2016).

13. Silva, A. et al. Output-driven feedback system control platform optimizes combinatorial therapy of tuberculosis using a macrophage cell culture model. PNAS 113, 2172–2179 (2016).

14. Cokol, M., Kuru, N., Bicak, E., Larkins-Ford, J. & Aldridge, B. B. Efficient measurement and factorization of high-order drug interactions in Mycobacterium tuberculosis. Sci. Adv. 3, e1701881 (2017).

15. Yeh, P., Tschumi, A. I. & Kishony, R. Functional classification of drugs by properties of their pairwise interactions. Nat. Genet. 38, 489–494 (2006).

16. Cicchese, J. M., Pienaar, E., Kirschner, D. E. & Linderman, J. J. Applying Optimization Algorithms to Tuberculosis Antibiotic Treatment Regimens. Cell. Mol. Bioeng. 10, 523–535 (2017).

17. Fonseca, K. L., Rodrigues, P. N. S., Olsson, I. A. S. & Saraiva, M. Experimental study of tuberculosis: From animal models to complex cell systems and organoids. PLoS Pathog. 13, 1–13 (2017).

18. Lin, P. L. et al. Quantitative comparison of active and latent tuberculosis in the cynomolgus macaque model. Infect. Immun. 77, 4631–4642 (2009).

19. Dartois, V. The path of anti-tuberculosis drugs: from blood to lesions to mycobacterial cells. Nat. Rev. Microbiol. 12, 159–167 (2014).

20. Prideaux, B. et al. The association between sterilizing activity and drug distribution into tuberculosis lesions. Nat. Med. 21, 1223–7 (2015).

21. Sarathy, J. P. et al. Prediction of Drug Penetration in Tuberculosis Lesions. ACS Infect. Dis. 2, 552–563 (2016).

22. Pienaar, E. et al. A computational tool integrating host immunity with antibiotic dynamics to study tuberculosis treatment. J. Theor. Biol. 367, 166–179 (2015).

23. Sarathy, J. P. et al. Extreme drug tolerance of mycobacterium tuberculosis in Caseum. Antimicrob. Agents Chemother. 62, 1–11 (2018).

24. Bowness, R., Chaplain, M. A. J., Powathil, G. G. & Gillespie, S. H. Modelling the effects of bacterial cell state and spatial location on tuberculosis treatment: Insights from a hybrid multiscale cellular automaton model. J. Theor. Biol. 446, 87–100 (2018).

25. Segovia-Juarez, J. L., Ganguli, S. & Kirschner, D. Identifying control mechanisms of granuloma formation during M. tuberculosis infection using an agent-based model. J. Theor. Biol. 231, 357–376 (2004).

26. Cilfone, N. A., Perry, C. R., Kirschner, D. E. & Linderman, J. J. Multi-scale modeling predicts a balance of tumor necrosis factor-a and interleukin-10 controls the granuloma environment during Mycobacterium truberculosis infection. PLoS One 8, e68680 (2013).

27. Ray, J. C. J., Flynn, J. L. & Kirschner, D. E. Synergy between individual TNF-dependent functions determines granuloma performance for controlling Mycobacterium tuberculosis infection. J. Immunol. 812, 3706–37017 (2009).

28. Fallahi-Sichani, M., El-Kebir, M., Marino, S., Kirschner, D. E. & Linderman, J. J. Multiscale computational modeling reveals a critical role for TNF-α receptor 1 dynamics in tuberculosis granuloma formation. J. Immunol. 186, 3472–3483 (2011).

29. Pienaar, E. et al. Comparing efficacies of moxifloxacin, levofloxacin and gatifloxacin in tuberculosis granulomas using a multi-scale systems pharmacology approach. PLOS Comput. Biol. 13, (2017).

30. Pienaar, E., Dartois, V., Linderman, J. J. & Kirschner, D. E. In silico evaluation and exploration of antibiotic tuberculosis treatment regimens. BMC Syst. Biol. 9, 1–12 (2015).

31. Cicchese, J. M., Dartois, V., Kirschner, D. E. & Linderman, J. J. Both Pharmacokinetic Variability and Granuloma Heterogeneity Impact the Ability of the First-Line Antibiotics to Sterilize Tuberculosis Granulomas. Front. Pharmacol. 11, 1–15 (2020).

32. Strydom, N. et al. Tuberculosis drugs’ distribution and emergence of resistance in patient’s lung lesions : A mechanistic model and tool for regimen and dose optimization. PLoS Med. 16, (2019).

33. Aljayyoussi, G. et al. Pharmacokinetic-Pharmacodynamic modelling of intracellular Mycobacterium tuberculosis growth and kill rates is predictive of clinical treatment duration. Sci. Rep. 7, 1–11 (2017).

34. Pasipanodya, J. G. et al. Serum drug concentrations predictive of pulmonary tuberculosis outcomes. J. Infect. Dis. 208, 1464–1473 (2013).

35. Bonnett, L. J., Ken-dror, G., Koh, G. C. K. W. & Davies, G. R. Comparing the Efficacy of Drug Regimens for Pulmonary Tuberculosis: Meta-analysis of Endpoints in Early-Phase Clinical Trials. Clin. Infect. Dis. 65, 46–54 (2017).

36. Gillespie, S. H. et al. Four-Month Moxifloxacin-Based Regimens for Drug-Sensitive Tuberculosis. N. Engl. J. Med. 371, 1577–1587 (2014).

37. Jindani, A. et al. High-Dose Rifapentine with Moxifloxacin for Pulmonary Tuberculosis. N. Engl. J. Med. 371, 1599–1608 (2014).

38. Pranger, A. D., van der Werf, T. S., Kosterink, J. G. W. & Alffenaar, J. W. C. The Role of Fluoroquinolones in the Treatment of Tuberculosis in 2019. Drugs 79, 161–171 (2019).

39. Zimmerman, M. et al. Ethambutol Partitioning in Tuberculous Pulmonary Lesions Explains Its Clinical Efficacy. Antimicrob. Agents Chemother. 61, 1–12 (2017).

40. Cokol, M., Li, C. & Chandrasekaran, S. Chemogenomic model identifies synergistic drug combinations robust to the pathogen microenvironment. PLoS Comput. Biol. 14, 1–24 (2018).

41. Zimmermann, M. et al. Integration of Metabolomics and Transcriptomics Reveals a Complex Diet of Mycobacterium tuberculosis during. mSystems 2, 1–18 (2017).

42. Chandrasekaran, S. Predicting Drug Interactions From Chemogenomics Using INDIGO. in Systems Chemical Biology. Methods in Molecular Biology 219–231 (2019).

43. GranSim. Available at: http://malthus.micro.med.umich.edu/GranSim/. (Accessed: 21st August 2020)

44. Pienaar, E., Matern, W. M., Linderman, J. J., Bader, J. S. & Kirschner, D. E. Multiscale Model of Mycobacterium tuberculosis Infection Maps Metabolite and Gene Perturbations to Granuloma Sterilization. Infect. Immun. 84, 1650–1669 (2016).

45. Cilfone, N. A. et al. Computational modeling predicts IL-10 control of lesion sterilization by balancing early host immunity-mediated antimicrobial responses with caseation during mycobacterium tuberculosis infection. J. Immunol. 194, 664–77 (2015).

46. Jonsson, S. et al. Population pharmacokinetics of ethambutol in South African tuberculosis patients. Antimicrob. Agents Chemother. 55, 4230–4237 (2011).

47. Greco, W. R., Bravo, G. & Parsons, J. C. The Search for Synergy: A Critical Review from a Response Surface Perspective. Pharmacol. Rev. 47, 331–385 (1995).

48. Hall, M. J., Middleton, R. F. & Westmacott, D. The fractional inhibitory concentration (FIC) index as a measure of synergy. J. Antimicrob. Chemother. 11, 427–433 (1983).

49. Mckay, A. M. D., Beckman, R. J. & Conover, W. J. A Comparison of Three Methods for Selecting Values of Input Variables in the Analysis of Output from a Computer Code. Technometrics 21, 239–245 (1979).

50. Marino, S., Hogue, I. B., Ray, C. J. & Kirschner, D. E. A methodology for performing global uncertainty and sensitivity analysis in systems biology. J. Theor. Biol. 254, 178–196 (2008).

51. Renardy, M., Hult, C., Evans, S., Linderman, J. J. & Kirschner, D. E. Global sensitivity analysis of biological multiscale models. Curr. Opin. Biomed. Eng. 11, 109–116 (2019).

