## Supplemental Tables and Figures for "A multi-scale pipeline linking drug transcriptomics with pharmacokinetics predicts *in vivo* interactions of tuberculosis drugs"

Supplementary Table 1. List of regimens simulated and their corresponding fractional inhibitory concentrations (FIC). The antibiotics included and their abbreviations and standard doses are isoniazid (H, 5 mg/kg), rifampin (R, 10 mg/kg), ethambutol (E, 20 mg/kg), pyrazinamide (Z, 25 mg/kg), moxifloxacin (M, 7 mg/kg), and levofloxacin (L, 17 mg/kg). The regimen names give the single letter abbreviation, followed by the dose in mg/kg for that antibiotic. If no number is listed, the standard dose was used. The doses per week (dpw) are listed at the end of the regimen. If no doses per week is listed, doses were simulated as administered daily. The regimens used for validation of the model are labeled with an asterisk, corresponding to the regimens listed in Figure 4 and referenced from Bonnet *et al.* (2017) (Bonnett et al. 2017).

| Regimen | FIC |
|---|---|
| HRZE | 0.82 |
| HR | 1.35 |
| HZ | 0.97 |
| HE | 1.46 |
| RZ | 1.33 |
| ZE | 1.12 |
| RE | 0.74 |
| HRZ | 0.74 |
| HRE | 0.94 |
| HZE | 0.91 |
| RZE | 0.78 |
| HMZE | 1.05 |
| RMZE | 0.96 |
| HM | 2.01 |
| RM | 2.31 |
| MZ | 1.99 |
| ME | 1.8 |
| HRM | 1.08 |
| HMZ | 1.1 |
| HME | 1.19 |
| RMZ | 1.33 |
| RME | 0.99 |
| MZE | 1.19 |
| HRMZ | 0.92 |
| HRME | 0.93 |
| HL | 1.25 |
| RL | 1.53 |
| LZ | 1.47 |
| LE | 1.28 |
| HRL | 1.01 |
| HLZ | 1.11 |
| HLE | 1.15 |

| Regimen | FIC |
|---|---|
| RLZ | 1.22 |
| RLE | 1.12 |
| LZE | 1.28 |
| HRLZ | 0.8 |
| HRLE | 0.89 |
| HLZE | 1.01 |
| RLZE | 0.94 |
| H6E15* | 1.46 |
| H6E25* | 1.46 |
| R11E25* | 0.74 |
| R23.5E25* | 0.74 |
| H11R10* | 1.35 |
| H6R10E16* | 0.94 |
| H6R10E25* | 0.94 |
| H10R10E25* | 0.94 |
| R24Z64E90dpw1* | 0.78 |
| H16R12Z49dpw2* | 0.74 |
| H6R10Z27E16* | 0.82 |
| H5R10M7Z25dpw5* | 0.92 |
| R10M7Z25E15* | 0.96 |
| H14R10M9Z34dpw3* | 0.92 |
| H6E6* | 1.46 |
| H10E25* | 1.46 |
| R23.5E45dpw2* | 0.74 |
| R23.5E90dpw1* | 0.74 |
| H6R6E10* | 0.94 |
| H15R10E40dpw2* | 0.94 |
| H5R10Z25E17.5* | 0.82 |
| H6R9Z24E16dpw3* | 0.82 |
| H14R13Z47E29dpw3 * | 0.82 |

*Used for validation

Supplementary Table 2 Host immune parameters used with *GranSim* to generate the granuloma bio-repository. Timestep units represent 10-minute time steps in the agent-based simulation. The minimum and maximum values for parameter ranges used in sampling are listed. Parameters are based on previous *GranSim* studies (Cicchese et al. 2020; Pienaar et al. 2015).

| **Parameter Definition** | **Units** | **Min** | **Max** |
|---|---|---|---|
| # immune cell deaths causing compartment caseation | | 6 | 10 |
| Time to heal caseated compartment | Timesteps | 909 | 1365 |
| TNF threshold for causing immune cell apoptosis | Molecules | 690 | 1035 |
| Rate constant for TNF-induced apoptosis | 1/s | 1.36e-6 | 2.04e-6 |
| Minimum chemokine concentration to induce chemotaxis | Molecules | 0.27 | 0.41 |
| Maximum chemokine concentration to induce chemotaxis | Molecules | 392 | 588 |
| Initial density of macrophages | Fraction of grid compartments | 0.019 | 0.029 |
| Time between resting macrophage movements | Timesteps | 4 | 6 |
| Time between active macrophage movements | Timesteps | 15 | 23 |
| Time between infected macrophage movements | Timesteps | 169 | 255 |
| TNF threshold to induce NFkB activation | Molecules | 42.8 | 64.1 |
| Rate constant for NFkB activation | 1/s | 6.77e-6 | 1.01e-5 |
| Probability resting macrophage kills extracellular Mtb | | 0.0738 | 0.111 |
| Killing probability adjustment for resting macrophages with NFkB activation | | 0.129 | 0.194 |

| | | | |
|---|---|---|---|
| # bacteria to cause NFkB activation | | 236 | 354 |
| # bacteria for macrophage to become chronically infected | | 12 | 18 |
| # bacteria to cause macrophage to burst | | 19 | 29 |
| # bacteria activated macrophage can phagocytose | | 3 | 5 |
| Probability activated macrophage will will heal a caseated compartment | | 0.00459 | 0.00687 |
| Probability a T-cell will move to same compartment as a macrophage | | 0.0367 | 0.0550 |
| Probability IFNγ producing T-cell induces Fas/FasL apoptosis | | 0.0293 | 0.0439 |
| Probability IFNγ producing T-cell also produces TNF | | 0.0514 | 0.0770 |
| Probability cytotoxic T-cell kills macrophage | | 0.00505 | 0.0121 |
| Probability cytotoxic T-cell kills a macrophage and all its intracellular bacteria | | 0.619 | 0.928 |
| Probability regulatory T-cell deactivates macrophage | | 0.00584 | 0.00876 |
| Time when T-cell recruitment begins | Timesteps | 3225 | 4722 |
| Time delay after T-cell recruitment begins until maximal recruitment rate | Timesteps | 650 | 976 |
| Macrophage maximal recruitment probability | | 0.0241 | 0.0361 |
| Macrophage threshold for recruitment by chemokines | Molecules | 0.641 | 0.960 |
| Macrophage threshold for recruitment by TNF | Molecules | 0.00859 | 0.0129 |

| | | | |
|---|---|---|---|
| Macrophage half saturation for recruitment by TNF | Molecules | 1.22 | 1.82 |
| Macrophage half saturation for recruitment by chemokine | Molecules | 1.68 | 2.52 |
| IFNγ producing T-cell maximal recruitment probability | | 0.0484 | 0.0726 |
| IFNγ producing T-cell threshold for recruitment by chemokine | Molecules | 0.0535 | 0.0802 |
| IFNγ producing T-cell threshold for recruitment by TNF | Molecules | 1.01 | 1.51 |
| IFNγ producing T-cell half saturation for recruitment by TNF | Molecules | 1.22 | 1.82 |
| IFNγ producing T-cell half saturation for recruitment by chemokine | Molecules | 1.64 | 2.46 |
| Probability a IFNγ producing T-cell is cognate | | 0.0437 | 0.0655 |
| Cytotoxic T-cell maximal recruitment probability | | 0.0370 | 0.0554 |
| Cytotoxic T-cell threshold for recruitment by chemokine | Molecules | 3.55 | 5.32 |
| Cytotoxic T-cell threshold for recruitment by TNF | Molecules | 0.920 | 1.38 |
| Cytotoxic T-cell half saturation for recruitment by TNF | Molecules | 0.715 | 1.07 |
| Cytotoxic T-cell half saturation for recruitment by chemokine | Molecules | 5.24 | 7.86 |
| Probability a cytotoxic T-cell is cognate | | 0.0414 | 0.0620 |
| Regulatory T-cell maximal recruitment probability | | 0.0246 | 0.0369 |
| Regulatory T-cell threshold for recruitment by chemokine | Molecules | 2.03 | 3.04 |

| | | | |
|---|---|---|---|
| Regulatory T-cell threshold for recruitment by TNF | Molecules | 1.65 | 2.47 |
| Regulatory T-cell half saturation for recruitment by TNF | Molecules | 2.00 | 3.00 |
| Regulatory T-cell half saturation for recruitment by chemokine | Molecules | 1.23 | 1.84 |
| Probability a regulatory T-cell is cognate | | 0.0400 | 0.0600 |

Supplementary Table 3. For four regimens (HRZE, RE, HE and RM), the table shows the PRCC values relating the plasma PK parameters to the predicted iDIS for non-replicating Mtb during the first dose of treatment. All PRCC values reported are significant with p<0.01 and NS designates that the parameter was not significant. The parameters for each antibiotic listed include the absorption rate constant (kAbs), the intercompartmental clearance (Q), the volume of distribution for the plasma (central) compartment (Vol. Dist. Cent.), the volume of distribution for the peripheral compartment (Vol. Dist. Periph.), and the clearance rate constant (CL).

| | INH | | | | | RIF | | | | | EMB | | | | | PZA | | | | | MXF | | | | |
|---|---|---|---|---|---|---|---|---|---|---|---|---|---|---|---|---|---|---|---|---|---|---|---|---|---|
| | kAbs | Q | Vol. Dist. Cent | Vol. Dist. Periph | CL | kAbs | Q | Vol. Dist. Cent | Vol. Dist. Periph | CL | kAbs | Q | Vol. Dist. Cent | Vol. Dist. Periph | CL | kAbs | Q | Vol. Dist. Cent | Vol. Dist. Periph | CL | kAbs | Q | Vol. Dist. Cent | Vol. Dist. Periph | CL |
| **HRZE** | NS | -0.66 | 0.34 | -0.18 | -0.20 | 0.28 | NS | NS | NS | 0.89 | 0.43 | NS | -0.26 | NS | -0.56 | NS | NS | 0.54 | NS | 0.21 | | | | | |
| **RE** | | | | | | NS | NS | NS | NS | 0.90 | 0.74 | -0.27 | -0.23 | -0.20 | -0.92 | | | | | | | | | | |
| **HE** | -0.31 | -0.87 | 0.67 | -0.33 | -0.39 | | | | | | -0.68 | 0.26 | 0.30 | 0.20 | 0.88 | | | | | | | | | | |
| **RM** | | | | | | NS | NS | NS | -0.26 | -0.83 | | | | | | | | | | | NS | -0.12 | 0.19 | 0.95 | 0.99 |

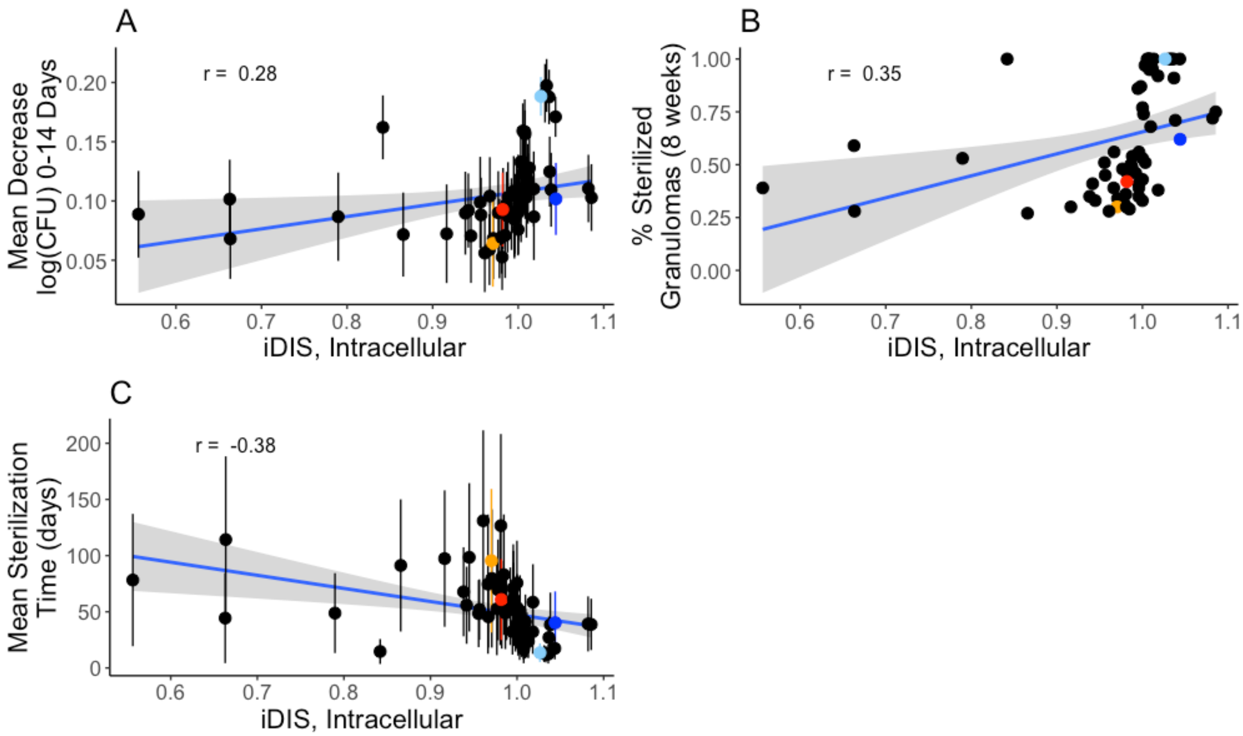


Supplementary Figure 1 Measures of regimen efficacy are correlated with interaction strength associated with intracellular replicating Mtb killing rate for 64 regimens. The mean decrease in log CFU (0-14) averaged over each 100 granulomas simulated for each regimen (A) and percentage of sterilized (negative) granulomas after eight weeks of treatment (B) are weakly positively correlated with the average interaction strength experienced by non-replicating Mtb during the first 24 hours of treatment with correlation coefficients of 0.28 and 0.35 respectively. Mean sterilization time for each regimen over 100 granulomas (C) is negatively correlated with the average interaction strength with a correlation coefficient of -0.38. Each point represents the regimen outcome measurement for a given regimen and error bars indicate +/- standard deviation from the sample of 100 granulomas simulated. The colored points correspond to the regimens HRZE (light blue), RE (dark blue), RM (red) and HE (orange).

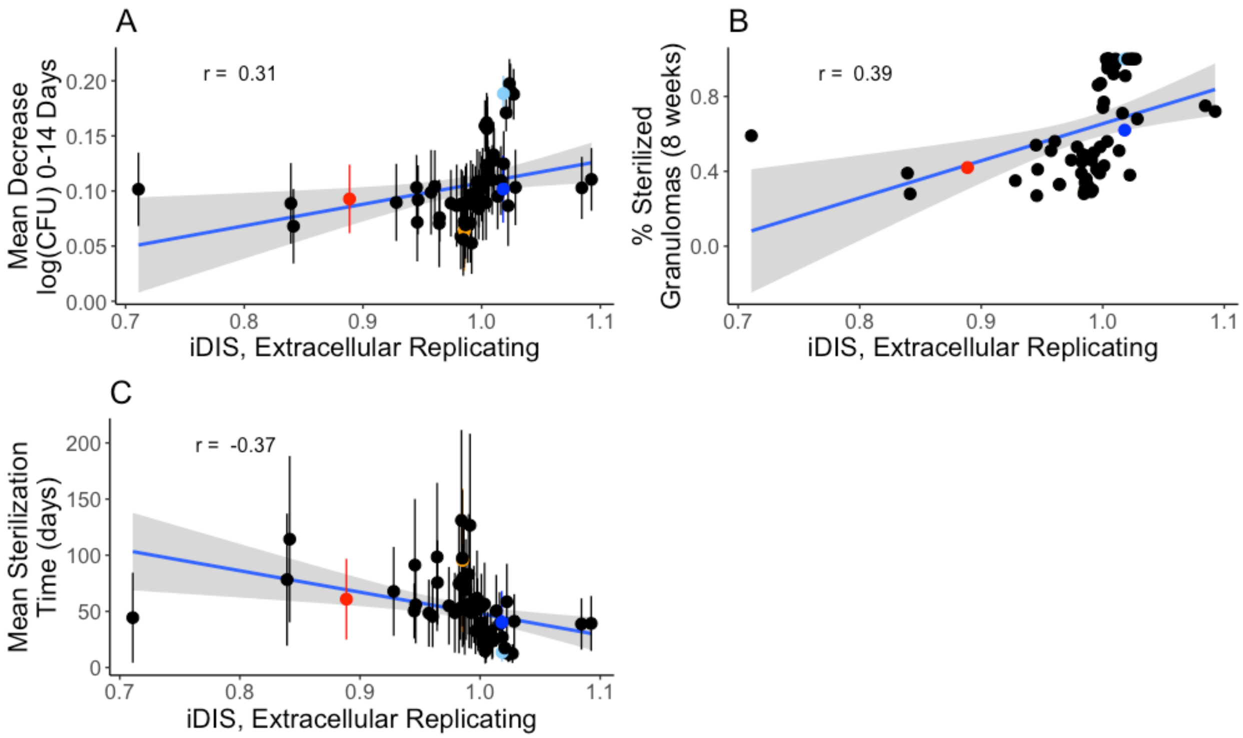


Supplementary Figure 2 Measures of regimen efficacy are correlated with interaction strength associated with extracellular replicating Mtb killing rate for 64 regimens. The mean decrease in log CFU (0-14) averaged over each 100 granulomas simulated for each regimen (A) and percentage of sterilized (negative) granulomas after eight weeks of treatment (B) are weakly positively correlated with the average interaction strength experienced by non-replicating Mtb during the first 24 hours of treatment with correlation coefficients of 0.28 and 0.35 respectively. Mean sterilization time for each regimen over 100 granulomas (C) is negatively correlated with the average interaction strength with a correlation coefficient of -0.38. Each point represents the regimen outcome measurement for a given regimen and error bars indicate +/- standard deviation from the sample of 100 granulomas simulated. The colored points correspond to the regimens HRZE (light blue), RE (dark blue), RM (red) and HE (orange).

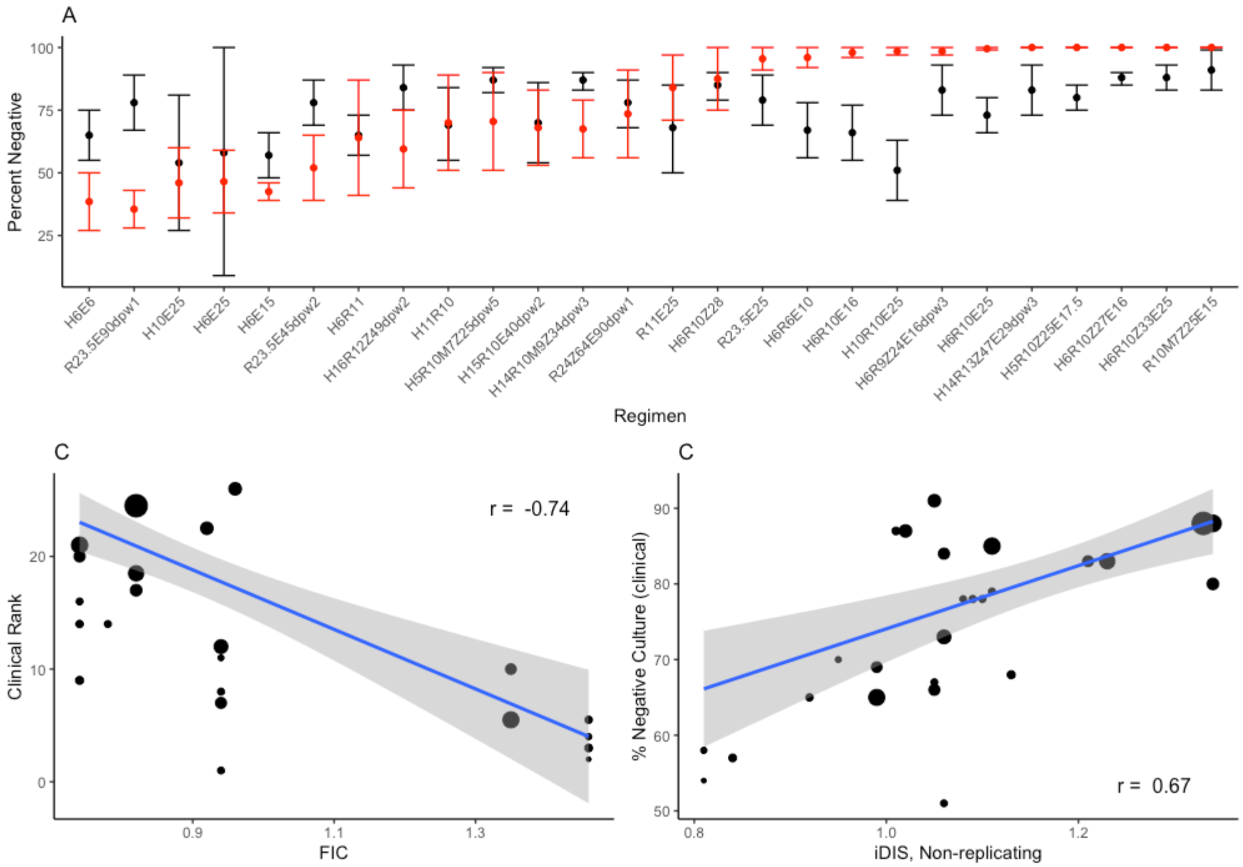


Supplementary Figure 3 Comparison of treatment simulations with clinical trial results for 26 different regimens compiled in Bonnet *et al.* (2017). The clinical regimen efficacy metric used is solid culture conversion following 8 weeks of therapy. We compare the confidence intervals (black) to the percent of simulated granulomas that sterilized (lower red bar) or had fewer than 10 CFU after 8 weeks of therapy (upper red bar, red dot average of error bars) for all 26 regimens (A). Regimens are abbreviated by the single antibiotic abbreviation, followed by the dose in mg/kg for that antibiotic, with the doses per week (dpw) listed at the end of the regimen abbreviation. FIC is negatively correlated with clinical rank with a weighted correlation of -0.74 (B), and iDIS is positively correlated with clinical rank with a weighted correlation of 0.67 (C). Each dot represents an individual regimen, its size is linearly scaled by the number of patients treated (B and C)

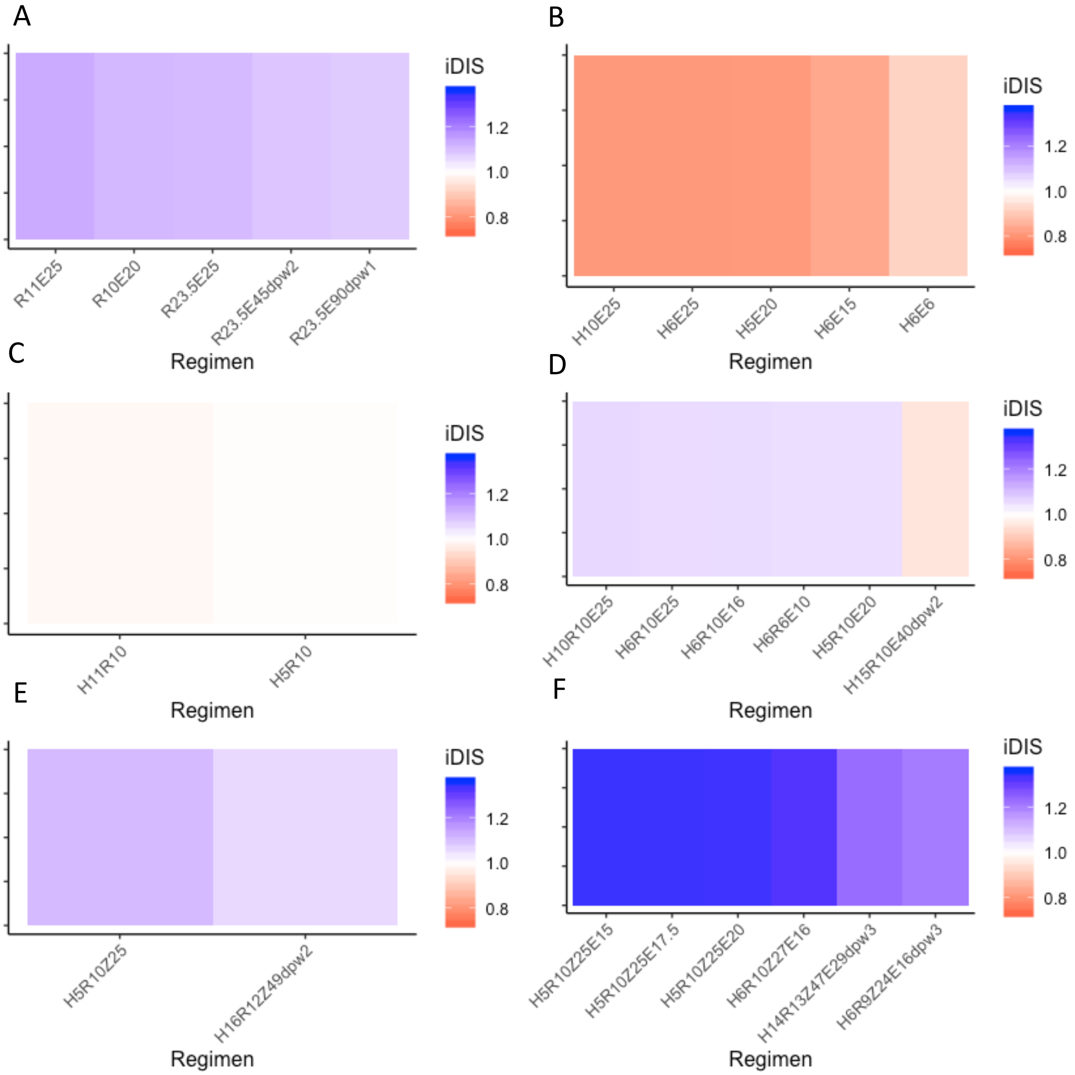


Supplementary Figure 4 Heat map of predicted iDIS value for different regimens of the same antibiotic combination. The list of regimens is ordered by decreasing predicted in vivo DIS for regimens involving the antibiotic combination RE (A), HE (B), HR (C), HRE (D), HRZ (E) and HRZE (F). For predicated IDIS blue represents synergy, white represents additivity, and red represents antagonism.

**References**

Bonnett, Laura J, Gie Ken-dror, Gavin C K W Koh, and Geraint R Davies. 2017. "Comparing the Efficacy of Drug Regimens for Pulmonary Tuberculosis: Meta-Analysis of Endpoints in Early-Phase Clinical Trials." *Clinical Infectious Diseases* 65(1): 46–54.

Cicchese, Joseph M., Véronique Dartois, Denise E. Kirschner, and Jennifer J. Linderman. 2020. "Both Pharmacokinetic Variability and Granuloma Heterogeneity Impact the Ability of the First-Line Antibiotics to Sterilize Tuberculosis Granulomas." *Frontiers in Pharmacology* 11(333): 1–15.

Pienaar, Elsje et al. 2015. "A Computational Tool Integrating Host Immunity with Antibiotic Dynamics to Study Tuberculosis Treatment." *Journal of Theoretical Biology* 367: 166–79. http://dx.doi.org/10.1016/j.jtbi.2014.11.021.
